# Heat stress drives increases in phytoplankton cell size in future oligotrophic oceans

**DOI:** 10.1101/2025.06.22.660956

**Authors:** Suzana G. Leles, Lara Breithaupt, Arianna Krinos, Harriet Alexander, Holly V. Moeller, Lana Flanjak, Charlotte Laufkotter, Elena Litchman, María Aranguren-Gassis, Naomi M. Levine

## Abstract

Warming and nutrient limitation are major stressors that affect primary production in the ocean, with cascading impacts on the food web. Yet, we lack a mechanistic understanding of how phytoplankton manage multiple stressors and the implications of these responses for phytoplankton biogeography. By combining theory, proteome allocation modeling, and climate projections, we identified two potential competing strategies for multi-stressor growth: (1) nutrient efficient smaller cells, or (2) heat-tolerant larger cells. We found that *Prochlorococcus* are more vulnerable in warmer, ‘heat-stressed’, tropical regions due to greater heat sensitivity and lower lipid storage capacity to buffer oxidative stress, indicating a potential ecological niche for larger phytoplankton with lower sensitivity to oxidative stress, such as *Synechococcus* and picoeukaryotes. Our findings advocate for the inclusion of phytoplankton heat-stress responses in global models to more accurately predict their ecological niches as the climate warms.

## INTRODUCTION

Future climate scenarios project an average increase of 1.5°C by 2040 in the global ocean^1^ with vast regions of the ocean warming to above 30°C by the end of the century^2^. Ocean warming is expected to promote stronger stratification, potentially decreasing the influx of nutrients from below the mixed layer^3^. Changing ocean conditions have already resulted in decreased primary production^4^ and shifts in species distributions^5^ and size spectra^6^. As phytoplankton drive ocean carbon cycling and fuel the ocean food web^7^, understanding how these microbes respond to changes in temperature and nutrient availability is critical for predicting ecosystem function as the climate changes^8–10^.

A key driver of microbial response to resource availability and temperature is cell size, more specifically surface area-to-volume ratio^11–14^. Under low nutrient concentrations, uptake is limited by molecular diffusion. Thus, a phytoplankter cell can increase nutrient uptake by increasing the number of transporters in the outer membrane^15^. As there is a limit to the number of transporters a membrane can support, cells maximize the nutrient flux per unit of cellular volume by decreasing cell size and thereby increasing their surface area-to-volume ratio^10,12,14^ (Figure 1). While warming temperatures often result in smaller cell sizes^16,17^, experimental^18–21^ and field^22^ studies have shown that phytoplankton increase their cell volume at critically high temperatures. In a previous study, we suggested that these larger cell sizes are a result of increased investment in energy metabolism and heat-mitigation, both of which require a lower surface area-to-volume ratio, i.e., larger cell sizes, to accommodate carbon storage and antioxidant machinery^23^ (Figure 1). This sets up a challenge for phytoplankton cells facing both nutrient and heat stress: invest in uptake which requires more transporters and higher surface area-to-volume ratios (smaller sizes), or invest in heat-stress mitigation strategies which require lower surface area-to-volume ratios (larger sizes) (Figure 1).

**Figure 1:**
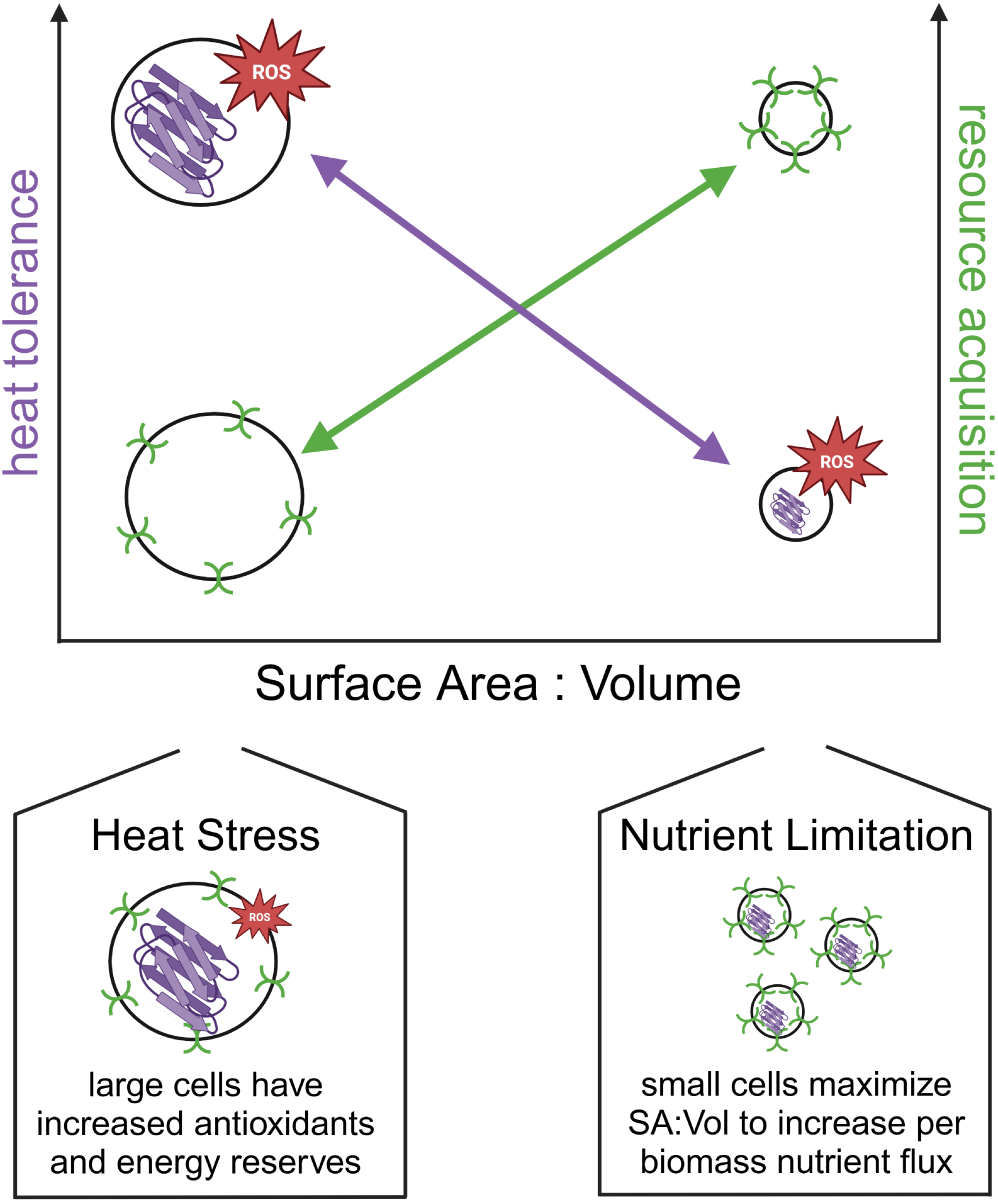
Predicted trade-off for phytoplankton under multi-stressor growth. Under nutrient limitation, phytoplankton need to invest more in transporters increasing their surface area-to-volume ratio, which requires smaller cell sizes. Under heat-stress, the cell must invest in heat-mitigation strategies decreasing their surface area-to-volume ratio to accommodate antioxidants and other proteomic investments which requires larger cell sizes. A trade-off between decreasing cell size to optimize nutrient uptake and increasing cell size to optimize heat-mitigation emerges under the multiple stressor scenario.

The global ocean has well known biogeographical distributions of phytoplankton^24^ with *Prochlo coccus* dominating the oligotrophic gyres and larger eukaryotic phytoplankton thriving in more nutrient-rich temperate and coastal regions^25^. Current models predict that, as the climate warms, the oligotrophic gyres will expand poleward resulting in an expanded ecological niche for *Prochlorococcus* ^26,27^. However, these predictions are based on a simplified representation of phytoplankton physiology that does not account for cellular strategies to mitigate heat stress and store carbon, both of which can promote increases in cell volume^19,21,22,28–31^. These predictions contrast with recent field data which demonstrates that *Prochlorococcus* abundance declines substantially in waters above 30 °C^32^. However, the mechanism for this constraint and why *Prochlorococcus* has not been able to evolve to manage elevated temperatures remains unknown. Constraining how cells respond to multi-stressor environments and how physiological responses interact to shape phytoplankton biogeography is critical for robustly predicting future ocean ecosystems.

Here, we investigate underlying mechanisms behind phytoplankton responses to co-occuring heat-stress and nutrient limitation using a proteome allocation model. We demonstrate that the thermal optimum for cells (*T_opt_*) can shift to either warmer or cooler temperatures under nitrogen limitation and that this shift is modulated by the heat-mitigation capacity of the cell. We then expand our theoretical predictions by calibrating the model to data for relevant phytoplankton functional types and simulate optimal growth strategies across a latitudinal transect under current and future projections of temperature and nitrate concentrations in the ocean. While we observe that the niche of cyanobacteria expands poleward, we also find that *Prochlorococcus* are more vulnerable in the warmer tropical seas due to their low heat-mitigation capacity and small cell sizes. We thus identify a critical new niche in warm tropical seas where phytoplankton with larger cell sizes, such as *Synechococcus* or even small eukaryotes, might thrive due to their larger flexibility in cell size and thermal tolerance. Our results suggest that representing the interaction between heat-stress and nutrient limitation in climate models is paramount for predicting species extinction and succession.

## RESULTS

### The proteome allocation model

We used a generic phytoplankton proteome allocation model to understand the molecular mechanisms behind changes in *T_opt_* and cell size under multi-stressor growth (Figure 1). Here, we expand the model of^23^ to simulate a range of phytoplankton phenotypes under the multiple stressor condition of heat-stress and resource limitation. The model can be generalized to all single-celled phytoplankton (prokaryotes and eukaryotes), but in its current form does not capture flexible metabolic strategies such as osmotrophy and mixotrophy. Given that nitrogen is often the most limiting nutrient within oligotrophic gyres^33^, we investigated phytoplankton responses to nitrogen limitation. However, the model dynamics of resource limitation are broadly applicable to other limiting nutrients (e.g., phosphorus). We then validated the model against thermal growth curve data and proteomic datasets for a range of phytoplankton species and use the model to simulate phytoplankton growth along a latitudinal gradient under current and future ocean conditions (see Methods).

Our proteome model consists of a system of ordinary differential equations defined as a constrained optimization problem (see Supplemental Methods for complete list of model equations). The model predicts cellular traits and physiological states by maximizing the steady-state growth rate of a phytoplankton cell under specific environmental conditions (i.e., irradiance, external inorganic nitrogen and carbon concentrations, and temperature). Specifically, the model simulates cellular resource allocation by optimizing the relative proteome investment across metabolic functions (e.g., nutrient uptake, biosynthesis, respiration), intracellular macromolecular concentrations (e.g., proteins, carbon storage), and the surface area-to-volume ratio of the phytoplankton cell (Table S1). The model resolves carbon and nitrogen metabolisms, including transport of nitrogen into the cell and synthesis of all critical cellular components. Parameter values are given in the Supplemental Methods (Tables S2 and S3).

Temperature accelerates all enzymatic reactions in the model (Equation 3, Methods; Figure S1) except photochemical reactions, which we assumed to be temperature independent^8,23^. Nitrogen transporters have a lower thermal dependency than other reactions in the model based on previous empirical^34–37^ and modeling^23^ studies. All other processes are assumed to have the same thermal dependency (defined by the parameter *E_a_*; Figure S1). The emergent thermal sensitivities between phototrophic and heterotrophic processes in our model are consistent with previous experimental and theoretical results^8,23^. Warming also results in protein damage in the model, thus mechanistically representing the decay in growth rates as temperatures increase beyond the optimum. To minimize heat-damage, the model cell can invest in repair pathways and in heat-mitigation strategies, such as producing antioxidant enzymes and/or storing fatty acids^38–40^. Heat-mitigation capacity is controlled by the model parameter *α*, with higher values enabling the cell to build more antioxidant machinery which, in turn, allows the cell to be more efficient at mitigating oxidative stress (Equation 4, Methods). While high *α* values provide protection against oxidative stress, this strategy comes with a trade-off of requiring intracellular space and proteomic investment that divests resources away from nutrient acquisition and/or growth.

We solve the proteome model over a range of temperatures and nitrogen concentrations, allowing thermal growth curves to emerge as a function of molecular constraints. Key to our analysis is that we are able to evaluate the underlying physiological and molecular mechanisms driving changes in thermal traits (e.g., *T_opt_*) and cell size. The optimal surface area-to-volume ratio of the cell is solved by the model as a function of surface area required for transporters in the cell membrane and cellular volume required for intracellular machinery, such as photosystems, nucleus, and antioxidants. Therefore, our model offers a platform to test, mechanistically, the cell size trade-off that phytoplankton face under the combined effects of heat-stress and nitrogen limitation (Figure 1).

### Understanding growth trade-offs under multi-stressor conditions

To understand the fundamental trade-offs associated with multi-stressor growth, we simulated phytoplankton cells with varying enzymatic thermal dependencies (*E_a_*) and heat-damage mitigation ability (*α*) and analyzed the emergent thermal growth responses (see Sensitivity Analyses, Methods). For this analysis, the cells were otherwise equal (i.e., all other parameters were equal). The model captured the expected trade-offs where warm-adapted cells (higher *E_a_*) have faster growth at higher temperatures but slower growth at lower temperatures (Figure 2A) and cells with higher heat-mitigation capacity (higher *α*) survive at higher temperatures at the cost of lower maximum growth rates (Figure 2B).

**Figure 2:**
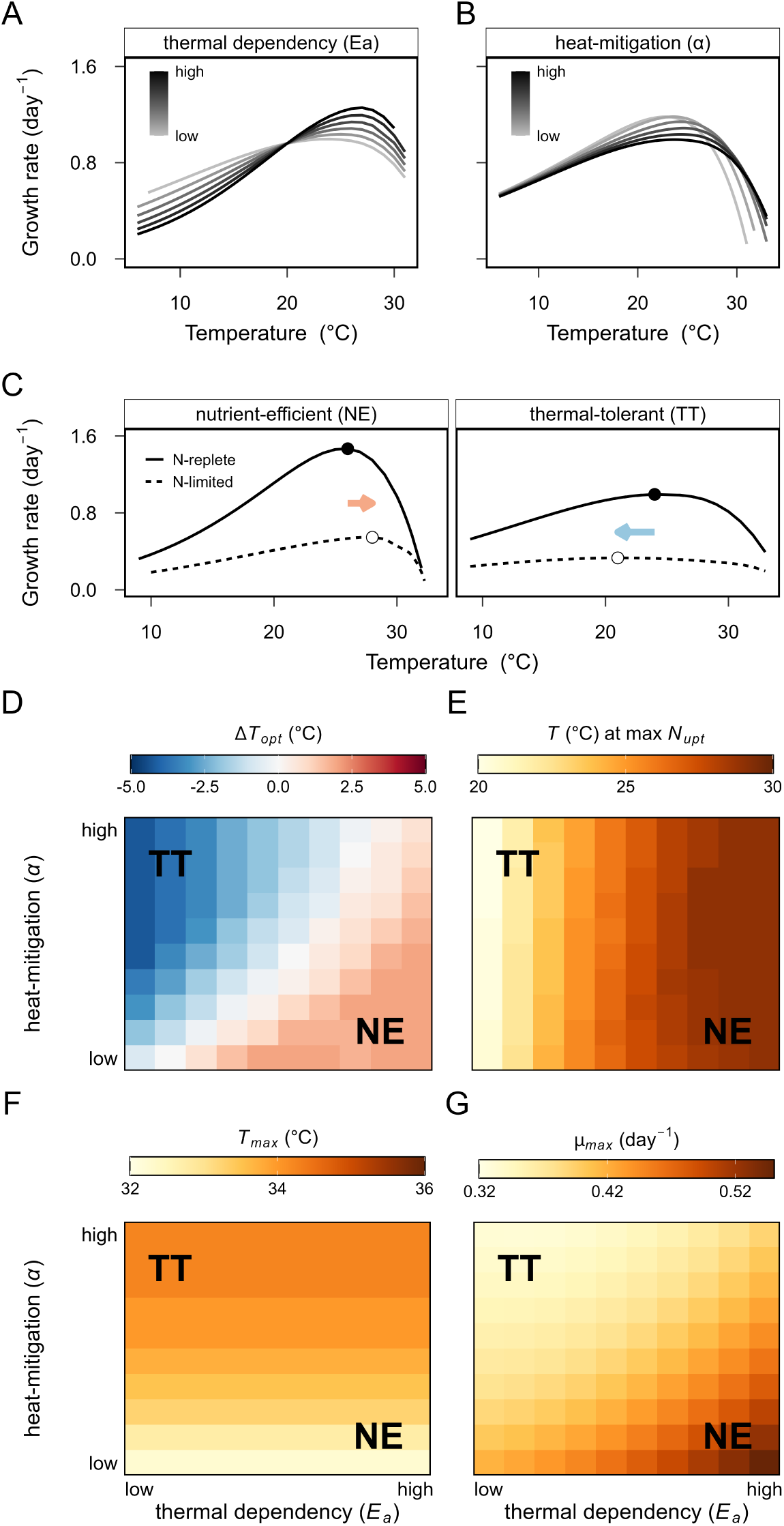
Growth trade-offs as a function of thermal dependency (*Ea*) and heat-mitigation capacity (*α*). Thermal growth curves under nitrogen-replete conditions are given for (A) cells with variable *E_a_* and (B) cells with variable *α*. (C) Thermal growth curves indicate shifts in *T_opt_* (open circles) from N-replete to N-limited conditions as a result of different metabolic strategies. (D) The change in *T_opt_* from N-replete to N-limited conditions (Δ *T_opt_*) showing increases (positive and red values) and decreases (negative and blue values) in *T_opt_*. (E) Temperature (*T*) where nitrogen uptake rates (*N_upt_*) are maximized under N-limited. (F) Maximal growth temperatures (*T_max_*) under N-limited. (G) Maximal growth rates (*µ_max_*) under N-limited. The two competing strategies are highlighted: nutrient-efficient (NE) and thermal-tolerant (TT).

The model predicts both increases and decreases in *T_opt_* under nitrogen limitation (Figures 2C and D), consistent with experimental results^41–43^. The model suggests that warm-adapted cells (higher *E_a_*) are more likely to increase their *T_opt_* under nitrogen limitation (Figure 2D), allowing cells to maximize nitrogen uptake rates at higher temperatures (Figure 2E). Higher heat-mitigation capacity allows the cell to survive at higher maximum temperatures (Figure 2F) but it also results in a decrease in *T_opt_* and lower maximum growth rates under nutrient stress (Figures 2D and G). Thus, two competing strategies for coping with multi-stressor conditions emerge from the model results: ‘nutrient-efficient’ or ‘thermal-tolerant’ (Figure 2C).

The ‘nutrient-efficient’ strategy couples higher *E_a_* values with lower *α* values. Higher *E_a_* values allow cells to invest primarily in growth (ribosomes). In turn, lower *α* values results in less investment in heat-mitigation strategies which allows for smaller cell sizes and more efficient nutrient uptake. This comes with the trade-off that cells are more susceptible to heat-damage and so have lower maximum growth temperatures. Alternatively, the ‘thermal-tolerant’ strategy cells invest heavily in heat mitigation (higher *α*). This results in increased proteomic investment in heat mitigation proteins which diverts resources away from growth (ribosomes) and results in lower maximum growth rates (Figure 2G). This strategy also requires more antioxidant machinery that results in larger cells. However, this investment allows for a higher maximum critical thermal limit (Figure 2F). Because the ‘thermal-tolerant’ strategy requires larger cell sizes, cells cannot be both thermally tolerant and nutrient efficient. Our model predicts that cells which combine low *E_a_* and low *α* would be maladapted with a lower maximal survival temperature under nitrogen limitation, with no growth advantage, in agreement with the “metabolic meltdown” theory^44^.

### Predicting phytoplankton optimal growth strategies in a changing ocean

We investigate the implications of different growth strategies in a changing ocean using four contrasting phytoplankton functional types: *Prochlorococcus*, *Synechococcus*, warm-adapted diatom, and cold-adapted diatom. Cyanobacteria are small and optimized for high nutrient use efficiency^45^, while diatoms have substantially larger cell sizes and carry large numbers of genes for mitigating temperature and oxidative stress^46–48^. However, differences are also observed within these two groups. Among the two cyanobacteria, *Synechococcus* have slightly larger cell sizes and are less sensitive to oxidative stress than *Prochlorococcus* ^49^. Between the two diatoms, the warm-adapted type has the potential for faster maximum growth than the cold-adapted diatom^50^. Diatoms were selected as a representative bloom forming eukaryotic phytoplankton due to their abundance throughout the oceans, the critical role they play in marine ecosystems, and the availability of data to validate the model. The dichotomy between cyanobacteria and diatoms in our model also mirrors the way phytoplankton diversity is typically simplified in biogeochemical models (see Discussion)^51^.

We parameterized the proteome model to represent these four strategies based on differences in nutrient uptake efficiency, thermal traits, and heat-mitigation capacity (Table S3, Methods). We validated our model phenotypes against thermal growth data from 167 species obtained from Anderson et al. ^50^. The maximum growth rate and slope of the thermal curves for our model phenotypes (Figure S2) show good agreement with observed values (Figure 3A). In addition, the emergent cell sizes in the model were consistent with the expected differences between these two groups (Figures 3B and S3). Because we focused on the impact of co-occurring temperature and nutrient stress, we chose to represent pico- and nano-sized diatoms since they are the direct competitors with cyanobacteria in the nutrient-deplete oligotrophic gyres, which are projected to undergo significant warming in the next century^2^. However, we demonstrate that the model can also simulate larger phytoplankton (Figure S4). We also qualitatively validated modeled shifts in proteome investment under heat or nutrient stress using proteomic datasets, based on the up- or down-regulation of specific pathways (Tables S4 and S5, see Methods). Unfortunately, no proteomic datasets were available to validate predictions under combined heat and nutrient stress.

**Figure 3:**
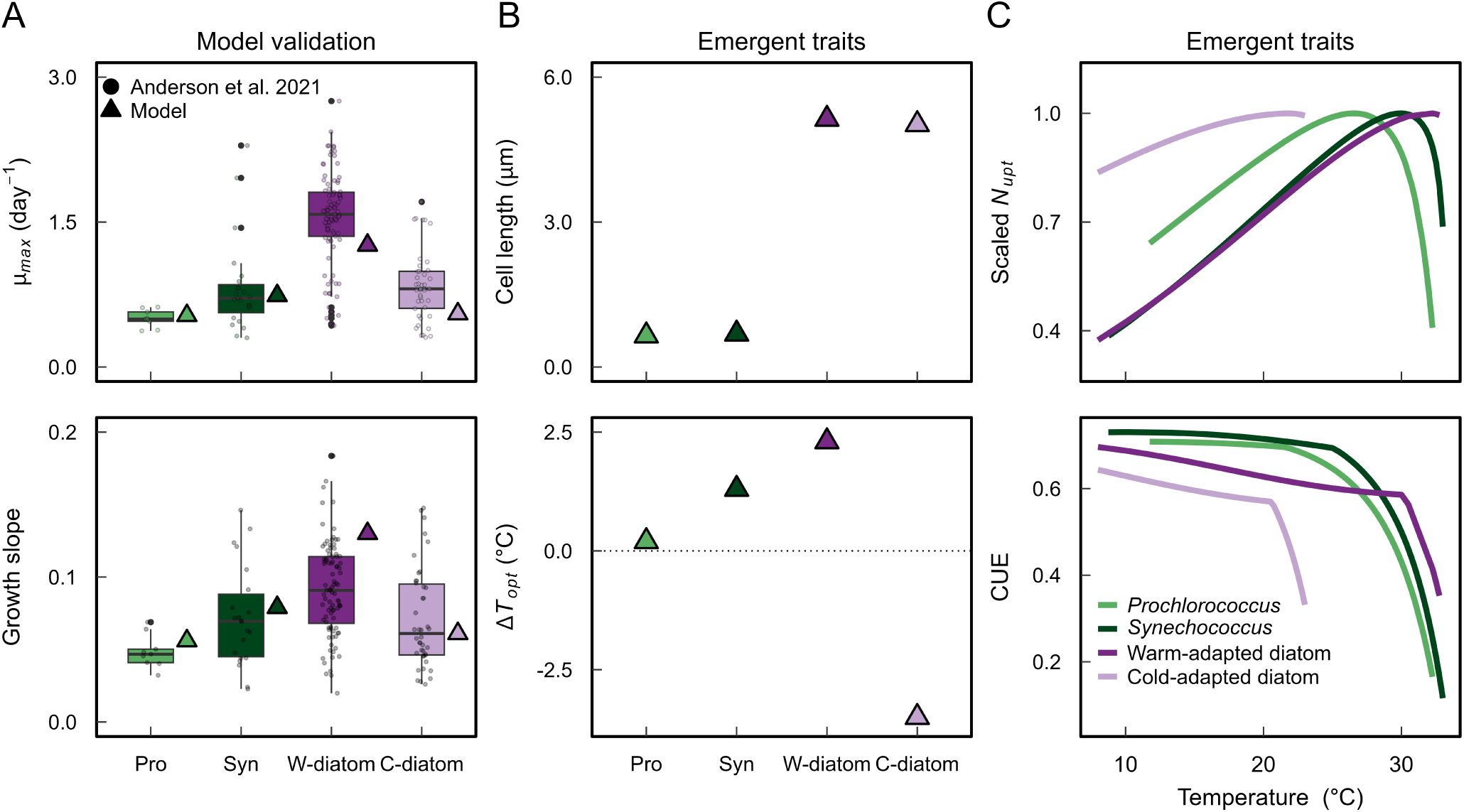
Emergent traits and trade-offs for different phytoplankton functional types. (A) Model validation against empirical thermal curve data from^50^ for the cyanobacteria *Prochlorococcus* (n = 9) and *Synechococcus* (n = 23) as well as cold-adapted (n = 42) and warm-adapted (n = 93) diatoms under nitrogen replete conditions. Maximum growth rates (*µ_max_*) and the slope at which growth rates increase as a function of temperature are shown. (B) Emergent model cell length at *T_opt_* under nitrogen limitation (see Figure S3 for N-replete conditions) and the shift in *T_opt_* from N-limited to N-replete conditions. (C) Model nitrogen uptake rates (*N_upt_*) scaled by maximum observed rate and Carbon Use Efficiencies (CUE) under nitrogen limitation. Carbon use efficiency is defined as 1 - respiration/photosynthesis. See Figures S2 and S5 for full thermal curves and proteome investments in both nutrient regimes.

Under heat stress, all phenotypes divested resources away from nitrogen transporters to invest in heat-mitigation strategies and glycolysis to support increased respiration demands (Figure S5). This trade-off is supported by observed proteomic shifts in the marine diatom *Chaetoceros simplex* ^52^, where populations down-regulated the investment in nitrogen transporters when grown at temperatures that induced heat-stress, while up-regulating the investment in chaperones, ROS scavenging, heat shock proteins, and glycolysis (Table S4). Under nitrogen stress, all modeled phenotypes maximized the specific affinity for nitrogen uptake by decreasing cell size (Figure S5). Cells also divested resources away from photosystems, rubisco, ribosomes, and respiration under nitrogen limitation, while investing more in transporters (Figure S5). These metabolic shifts are overall consistent with empirical observations for both diatoms and cyanobacteria (summarized in Table S5).

Under multi-stressor growth, the ‘nutrient-efficient’ strategy allowed *Prochlorococcus* to achieve the fastest maximum growth rates (Figure S2). However, at critically high temperatures, the ‘thermal-tolerant’ strategy allowed *Synechococcus* and the warm-adapted diatom to achieve higher carbon and nitrogen use efficiency (Figure 3C), resulting in better survival at higher maximum temperatures (Figure S2). In turn, the lower thermal dependency of the cold-adapted diatom resulted in the slowest growth rates under multi-stressor conditions.

We then use these four phenotypes to investigate optimal growth strategies across a latitudinal transect in the Atlantic Ocean under present day (1980-2000) and future (2080-2100) conditions (see Methods; Figure 4). For this analysis, we use temperature and nitrate concentrations for these two time periods from the CESM-WACCM model (Figure 4B). The proteome model captured present-day biogeographical distributions. Specifically, we observed the expected distributions of cold-adapted diatoms at the poles and warm-adapted diatoms in the temperate ocean due to their different thermal dependencies (Figures 4C and D). The cyanobacteria showed the highest growth potential in the oligotrophic gyres as expected (Figure S6), especially in the ultra-oligotrophic South Atlantic, due to their high specific nutrient affinity (Figures 4C and D). The warm-adapted diatom dominated again near the Equator due to increases in nitrate concentrations, except in the latitudinal range 0-7°N where nitrate concentrations were minimal and temperatures were maximal (Figures 4C and D). We would like to note that our model only predicts potential growth advantage since we did not simulate ecological interactions and physical dynamics (see Discussion).

**Figure 4:**
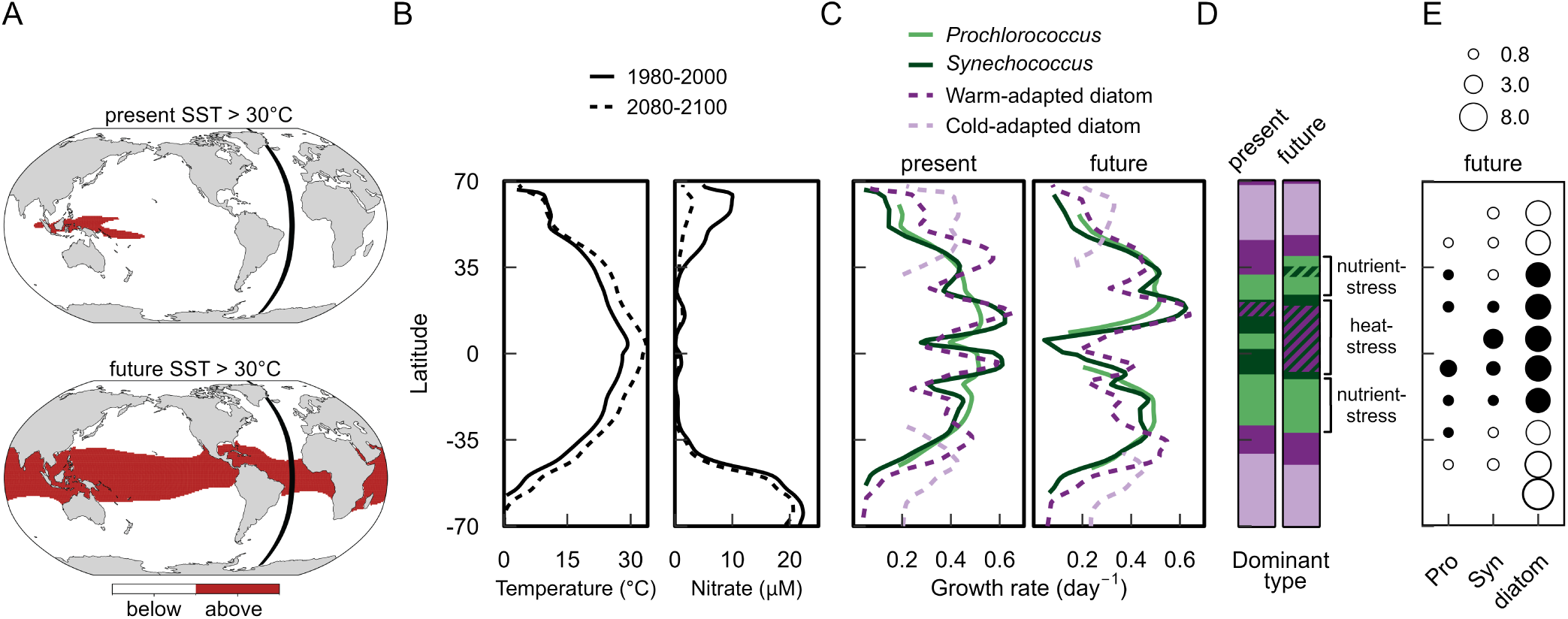
Phytoplankton optimal growth strategies across a latitudinal gradient. Future projections from the CESM-WACCM model indicate that 34.5% of global surface waters will experience temperatures above 30°C (A). Simulations were run using current (1980-2000) and future (2080-2100) temperature and nitrate concentrations along a latitudinal transect in the Atlantic Ocean (B). Also shown are the optimal growth rates (C), dominant functional type by latitude (D), and optimal cell diameter (*µm*) under the future climate scenario (E), where white fill and black fill shows decreases and increases in cell size from present to future, respectively. See Figure S9 for modeled proteome investments.

Under the future climate scenario, the model predicts a drastic increase globally in surface waters experiencing temperatures above 30°C from only 2.7% under present day to 34.5% by the end of the century (Figures 4A and S7). Temperatures along the transect also warmed significantly with average increase of 2.8 °C (maximum of 4.76 °C) and nitrate concentrations were on average 33% lower (maximum of 86%). The poleward expansion of the oligotrophic gyres, where decreases in nitrate concentrations were greater than the increases in temperature, resulted in the poleward expansion of cyanobacteria due to their higher nutrient affinity, i.e. ‘nutrient-efficient’ strategy (Figure 4D). The cold-adapted diatom phenotype still achieved the highest growth rates in the poles, however, the potential ecological niche for this group was narrower in the future projection due to the higher temperatures (Figure 4D).

The proteome model results also suggest a new niche where the projected increases in temperature will have a larger impact in governing phenotype shift than the projected change in nutrients (‘heat-stressed regions’, Figure 4D). This niche occurred in regions of the ocean that are severely nutrient-limited under present-day conditions and were predicted to experience significant warming (e.g., 12°S to 22°N). We hypothesize that the ‘thermal-tolerant’ strategy rather then the ‘nutrient-efficient’ strategy will have a competitive advantage in these regions, resulting in increases in cell size corresponding to the need to combat oxidative damage and meet dark respiration demands (Figure 4E). Thus, the *Synechococcus* and the warm-adapted diatom functional types are predicted to fair better in these ‘heat-stressed’ regions than *Prochlorococcus*. This finding holds even when assuming the same heat-mitigation abilities between the functional types (Figure S8), because *Prochlorococcus* has a lower thermal dependency than *Synechococcus* (Table S3), implying that thermal stress develops at lower temperatures. While the specific latitudinal range over which these changes would occur depend on the CESM model projections for changes in temperature and nitrate concentrations, our hypothesized shift in ecological niches provide insight into how the phytoplankton community will respond to multi-stressor change.

The underlying metabolic shifts in response to future multi-stressor change varied across oceanographic regions and by phenotype (Figures S9 and S10). At the poles, all phenotypes invested more in transporters (Figure S9) and decreased their cell size (white filled symbols in Figure 4E) to maximize nitrogen use efficiency under future more nitrogen limited conditions. This result is consistent with empirical observations that nitrogen limitation plays a more important role than temperature in determining phytoplankton cell size^14^. In the tropics, all phenotypes divested away from transporters and invested more in photosystems due to critical high temperatures, which increased respiration demands and oxidative stress, resulting in lower growth rates relative to current conditions (Figures S9 and S10). This strategy required larger cell sizes relative to present conditions (black filled symbols in Figure 4E). While *Prochlorococcus* (‘nutrient-efficient’ strategy) increased cell size to accommodate intracellular energy storage compounds to manage increasingly high rates of heat-damage and respiration, *Synechococcus* and the warm-adapted diatom (‘thermal-tolerant’ strategy) increased cell size to both store carbon for dark respiration and accommodate intracellular machinery for combating oxidative damage, thus better managing levels of heat-stress. This difference explains why the growth potential of *Prochlorococcus* was lower in the tropics. Overall, *Prochlorococcus* must devote a larger fraction of the proteome to respiration relative to *Synechococcus* and diatoms due to their lower heat-mitigation capacity (Figure S10). As a result, our model suggests that cyanobacteria, in particular the streamlined ultra-oligotrophic ‘nutrient-efficient’ *Prochlorococcus*, will have lower metabolic flexibility to manage a warmer climate.

## DISCUSSION

By combining ecological theory, proteome allocation modeling, and future climate projections, we were able to identify proteomic and cell size constraints underlying phytoplankton response to multi-stressor change. Specifically, our approach allows us to differentiate between ‘hard constraints’ imposed by physics (e.g., space and energy limitations) and ‘flexible constraints’ due to biology (e.g., enzyme kinetics)^23^. Unlike ‘hard constraints’, ‘flexible constraints’ can be mitigated through adaptation. We identified two different strategies that allow phytoplankton to minimize the negative impacts of climate warming under nitrogen limitation: ‘nutrient-efficient’ and ‘thermal-tolerant’. Due to hard constraints on cells, these two strategies are mutually exclusive - a cell cannot be both maximally nutrient efficient and have a high thermal tolerance. The cyanobacteria *Prochlorococcus* is a key example of the nutrient-efficient strategy. This strategy has allowed this group to thrive in present day oligotrophic gyres and we predict that their niche will expand polewards under future conditions, consistent with previous studies^26,27^. However, we demonstrate that evolving to become maximally nutrient-efficient, as *Prochlorococcus* has done, comes at a cost of thermal-tolerance. We therefore predict that future warming will result in the expansion of a ‘heat-stressed’ niche in the tropics where ‘thermal-tolerant’ phytoplankton will gain an advantage over ‘nutrient-efficient’ phytoplankton, due to their higher cell size flexibility and ability to manage heat-damage.

Our results highlight that cell size constraints are an important driver of phenotype. All phenotypes were predicted to decrease cell size in the polar and temperate oceans in response to nutrient stress and increase cell size in the tropics due to temperature stress (Figure 4E). This increase in cell volume for our model diatoms is consistent with previous work which has shown that diatoms and other eukaryotes increase cell size when exposed to temperatures above their growth maxima^18–20,22^ and when deprived of nutrients during transient stress, due to an accumulation of carbohydrates^29,31^ and/or lipids^28^. Similarly, our model predicted that, in order to survive in the tropics, the cyanobacteria functional types would need to significantly increase their cell diameter (up to 4-fold for *Prochlorococcus* and 3-fold for *Synechococcus*) to store carbon and support high respiration costs. This is consistent with recent laboratory^20,21^ and field manipulative experiments by^30^, which found that cyanobacteria increased their cell diameter under warming conditions up to 2-fold in the Atlantic Ocean, and that increases in cell volume were associated with slow or negative growth rates.

Global climate models predict a drastic increase in ultra-warm oligotrophic waters^2^, from 2 to 34% by the end of the century (Figures 4A and S7). These projections are based on the average temperature and do not show additional effects of temporal variability such as heatwaves, which will further exacerbate the situation. The two closest analogies in the present day ocean for these future conditions are the South Pacific Gyre and the Red Sea. While *Prochlorococcus* is abundant and the key contributor to carbon fixation around the rims of the South Pacific Subtropical Gyre, carbon fixation within the gyre is dominated by small photosynthetic eukaryotes^53^. An analysis of a large dataset (800 billion cells) from the Pacific demonstrated that *Prochlorococcus* is rarely found in waters above 30°C^32^. Red Sea surface temperatures can reach record values of 33°C in the surface^54^ and nutrient concentrations can be as low as 0.1 *µM* ^55^. Field observations show that the larger, more thermally tolerant cyanobacteria *Synechococcus* can dominate over *Prochlorococcus* in the surface waters of the Red Sea^55,56^, especially in the warmest (*>* 30°C) waters of the southern Red Sea^57^. While not conclusive, this does suggest that the Red Sea strains of *Prochlorococcus* have not been able to evolve to be competitive in the high temperature (*>* 30°C) niche^21^. In contrast, Synechococcus was shown to be able to easily evolve to growth at elevated temperatures (*>* 30°C) by increasing cell size and chlorophyll per cell^20^. Thus, both our work and field and experimental evidence suggest that elevated temperatures in the equatorial and tropical regions may favor small eukaryotes and larger cyanobacteria relative to *Prochlorococcus*.

We hypothesize that the ability to manage oxidative stress will play a key role in the success of different phenotypes in the future oceans that experience high levels of warming. *Synechococcus* are less sensitive to oxidative stress than *Prochlorococcus* ^49^. Overall, eukaryotes are better at sensing the environment and controlling oxidative stress relative to prokaryotes^46^, which is probably related to the lack of a large number of antioxidant genes in the latter, especially for *Prochlorococcus* ^47^. Previous work suggested that *Prochlorococcus* may have lost many antioxidant genes because they are costly and, instead, rely on other microbes to degrade reactive oxygen species for them^58,59^. Our model did not account for the potential benefit of these “helpers”. Yet, relying on other microbes to perform this important function may pose a risk to the adaptive potential of *Prochlorococcus*, as the “helpers” will also rapidly evolve under changing ocean conditions with no guarantee that the current interactions will persist^60,61^. Finally, diatoms and other eukaryotes have been found to associate symbiotically with nitrogen-fixing cyanobacteria, representing yet another strategy that enables larger eukaryotic cells to thrive under nitrogen depleted conditions^62,63^. Exploring how these microbial interactions will shift under multi-stressor changes is an important avenue for future work.

A key challenge faced broadly by numerical models is how to represent the vast diversity of marine microbes. For this study, we used diatoms as our example eukaryotic phytoplankton type to facilitate validation against experimental datasets and to be consistent with the simplified set of phytoplankton functional groups typically used in global biogeochemical models^2^. As we do not incorporate key diatom specific metabolisms such as silicate frustule synthesis into the proteome model, these findings are broadly applicable across bloom forming eukaryotic phytoplankton (e.g., haptophytes). However, our model currently does not resolve mixed forms of metabolisms, such as mixotrophy, which could be highly advantageous under multi-stressor environments. A key area for future research, both modeling and laboratory, is to understand the different strategies for coping with high temperature growth and oxidative stress in eukaryotic phytoplankton. Similarly, we focus on the upper water column (upper 20 m) where temperature and nutrient stress are greatest. However, there are sharp gradients within the euphotic zone and key differences between high and low light adapted *Prochlorococcus*. An additional area for future work would be to explore how the environment at the deep-chlorophyll maximum is predicted to shift and the impact on the relative competitive advantage of different strategies.

Here, we highlight the key importance that heat stress can play in determining biogeographical distributions of phytoplankton. Moreover, we hypothesize that due to inherent space and energy trade-offs, there are two alternative strategies for dealing with multi-stressor growth: ‘nutrient-efficient’ and ‘thermal-tolerant’. Our work advocates for the inclusion of heat stress strategies, carbon storage, and changes in cell size into Earth System Models used for future climate predictions. The proteome-allocation model results elucidate the trade-offs related to different metabolic strategies thereby providing a road-map for how to represent these dynamics in large-scale models. This would allow for future studies to investigate the interactive effects of cellular-level responses to multi-stressor growth with populations dynamics and ecological interactions.

The creation of a new niche for larger phytoplankton in the equatorial and tropical ocean under warming conditions has the potential to reshape microbial interactions, food web structure, and nutrient cycling. While we focused on pico- and nano-sized cells, we show that larger eukaryotes (Figure S4) are subjected to similar trade-offs (Figures S11 and S12). Eukaryotic phytoplankton typically release more diverse and abundant organic compounds than cyanobacteria, fueling heterotrophic bacterial growth and enhancing nutrient remineralization^64^. In addition, larger phytoplankton cells can support larger zooplankton, enhancing trophic transfer efficiency to larger predators^65^. Larger phytoplankton also have higher cellular carbon content, produce sticky exopolymers that promote particle aggregation^64^, and are consumed by zooplankton that generate denser fecal pellets. Together, these traits could increase the potential for vertical carbon export to deeper waters, altering the stoichiometry and depth of nutrient recycling. Integrating the insights from cellular scale models into biogeochemical models provides a powerful tool for synthesizing experimental and field data on cellular responses and scaling-up to understand large-scale climate impacts.

## RESOURCE AVAILABILITY

### Lead contact

Requests for further information and resources should be directed to and will be fulfilled by the lead contact, Suzana G. Leles.

### Materials availability

This study did not generate new unique reagents.

### Data and code availability

The model was written in Julia^66^ and the code developed in Leles and Levine ^23^ is deposited on Zenodo (DOI: https://doi.org/10.5281/zenodo.8125732). The model code used to run the analyses described in the present study are available on GitHub (https://github.com/LevineLab/ProteomePhyto/tree/main) and will be added permanently to Zenodo after acceptance. The CESM2-WACCM model output used in this study is available via the Earth System Grid Federation (https://esgf-node.ipsl.upmc.fr/projects/esgf-ipsl/).

## ACKNOWLEDGMENTS

We acknowledge funding from the: Simons Foundation through a Simons Postdoctoral Fellowship in Marine Microbial Ecology to SGL (877215), National Science Foundation (NSF) CAREER OCE grant to NML (2044852), Swiss National Science Foundation grant to CL and LF (203448), Xunta de Galicia Postdoctoral Fellowship (2013, 2016) and University of Vigo Research Talent Retention Program (2019) to MAG. AIK was supported by the Simons Collaboration on Computational Biogeochemical Modeling of Marine Ecosystems award (549931).

## AUTHOR CONTRIBUTIONS

S.G.L. and N.M.L. conceptualized this study and wrote the initial manuscript. S.G.L. developed the model, run simulations, and performed data analyses. L.B. developed code to improve the optimization algorithm. A.K. developed code to perform the analyses with the empirically derived thermal growth data. L.F. provided CESM2-WACCM model output. M.A.G. analyzed the proteomic dataset. H.A., H.V.M., C.L., E.L. and other authors revised the initial draft contributing to the final version of this manuscript.

## STAR METHODS

### KEY RESOURCES TABLE

#### EXPERIMENTAL MODEL AND STUDY PARTICIPANT DETAILS

This study did not involve any new experiments and exclusively relies on computer simulations. Simulations were compared against empirical data previously published in the literature as described in the Key resources table.

### METHOD DETAILS

#### The model

We use a proteome allocation model modified from Leles and Levine ^23^ to predict the optimal metabolic strategy and cell size of a phytoplankton cell under the multi-stressor growth condition of heat and nutrient stress. We further integrate the model to empirical data from 167 species previously compiled by Anderson et al. ^50^ to model the optimal strategies of different phytoplankton functional types across a latitudinal transect under current and future ocean conditions. Our coarse-grained model is defined as a constrained optimization problem that maximizes the steady-state growth of the cell. Here, we give the general equations governing the model behavior and focus on the processes relevant for the present work. For the full model equations and list of optimized variables (Table S1) and parameter values (Tables S2 and S3), we refer the reader to the Supplemental Methods and the code available on GitHub (https://github.com/LevineLab/ProteomePhyto/tree/main).

The model assumes a cell that is growing exponentially given a certain temperature and nitrogen concentration. The macromolecular concentrations (*c*) can thus be obtained assuming that the sum of the syntheses rates (*v_synthesis_*) minus the sum of the degradation rates (*v_degradation_*) are balanced by the cellular growth rate, *µ*:

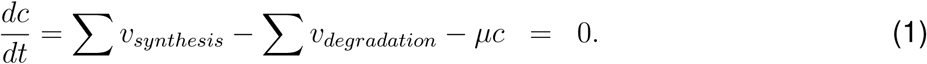

Each reaction rate catalyzed by a given protein *i* is described following a Michaelis-Menten type equation:

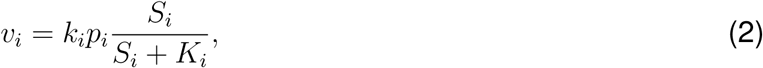

where *k_i_* is the maximum turnover rate per protein, *S_i_* is the substrate concentration, *K_i_* is the half-saturation constant, and *p_i_* is the concentration of protein *i*. *k_i_* is derived from literature values and corrected to a reference temperature of 20 °C. We assume all reaction rates in the cell increase with temperature following the Arrhenius equation, such that:

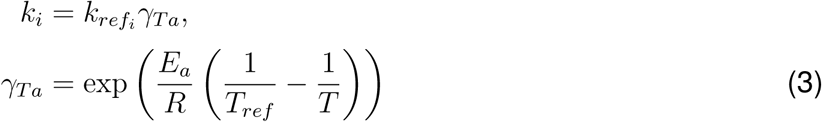

where 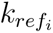 is the maximum turnover rate at a reference temperature *T_ref_* (in Kelvin) for a given protein *i*, *E_a_* is the thermal dependency of metabolic rates (eV), *R* is the Boltzmann’s constant (eV K^−1^) and *T* is the actual temperature (K). The only exception is the maximum rate of photosynthesis, which we assume to be independent of temperature^8^. All reactions were assumed to have the same *E_a_*, with the exception of nitrogen uptake, for which we halved the thermal dependency relative to other reactions in the model according to previous empirical^34–36^ and modeling^23^ studies.

Warming also results in protein damage *v_d_* which increases with increasing temperature, however, cells can invest in heat-mitigation strategies that reduce the impact of heat damage:

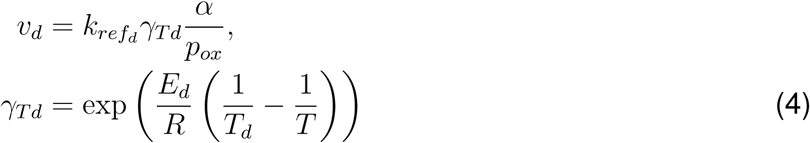

in which 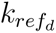 is the maximum rate of damage at a given temperature, *E_d_* is the thermal dependency of protein damage, *α* is the heat-mitigation capacity of the cell, and *p_ox_* is the pool of antioxidant machinery. *p_ox_* is constrained to be lower or equal to *α*. Thus, higher *α* enables the cell to build more antioxidant machinery *p_ox_*, which increases the efficiency of the cell in managing oxidative stress, but it comes at the cost of lower maximum growth rates (Figure 2B) due to higher intracellular space and proteomic costs required to build *p_ox_* that are ultimately divested away from growth. Our model can represent two different strategies in which a cell can mitigate heat-damage: storing lipids to sequester reactive oxygen species^39^ or producing antioxidant enzymes to decrease oxidative stress^40^. Here, we present the results for the model version in which a cell utilizes lipid storage. The results for antioxidant enzymes are quantitatively the same and are provided in the Supplemental Materials (Figure S13).

The model optimizes for the surface area-to-volume of the cell. We assume a spherical cell for simplicity and our model does not describe chains or filaments. The cell size is a trade-off between 1) surface area required for nitrogen transporters and 2) intracellular space required for intracellular machinery, including photosystems, carbon storage, nucleus, and antioxidants (see Supplementary Methods). The model can be applied to simulate a phytoplankton cell of any shape, but here we assume a spherical cell for simplicity.

#### Sensitivity analyses

Empirical observations show that phytoplankton species display diverse thermal growth curves^5,50^ and sensitivities to oxidative stress^46,67^. To capture such trait diversity, we varied two parameters in our simulations: the thermal dependency of metabolic reactions (*E_a_*) and the heat-mitigation capacity of the cell (*α*). We leveraged 167 phytoplankton thermal growth curves from previous compilations^5,50^ to select the range of *E_a_* values to test in our sensitivity analyses. Note that the value for *E_a_* in our model corresponds to the metabolic reaction and not necessarily the *E_a_* for growth rates, which is the commonly described parameter in the literature.

In turn, the value of *α* is poorly constrained in the literature. The range of *α* values used in our simulations was guided by a previous study, which found that phytoplankton sensitivity to oxidative stress can vary up to 10-fold between species^46^. This sensitivity was evidenced by elevated F_0_ and decreased oxygen production under H_2_O_2_ exposure, consistent with PSII photoinhibitory damage. Our model captures this differential sensitivity through the modeled concentrations of damaged photosystems (Figure S14). We further assumed that, the higher the maximum concentration of antioxidants *α*, the lower the energetic cost associated with protein damage *e_d_*; all remaining parameter values were held constant for these analyses (Table S2).

We performed simulations for 100 different cells based on a 10×10 grid combining different *E_a_* and *α* values. We solved the model across a wide range of temperatures for every 0.25 °C to capture the full thermal growth curves and derive the *T_opt_* for each phenotype from our simulations. We repeated this process for two different nutrient regimes to simulate nitrogen-replete (DIN = 20 *µ*M) and nitrogen-deplete (DIN = 0.1 *µ*M) conditions. In total, we simulated 200 thermal growth curves and found optimal solutions for approximately 10,000 combinations of temperature, DIN, *E_a_* and *α* (see Supplementary Methods).

#### Model comparison against proteomic data

We validated the heat-stress predictions from our model against data from an experimental evolution study that quantified the proteomic responses of the marine diatom *Chaetoceros simplex* ^52^. In the experiment, protein differential expression was compared between three different populations: ancestral population acclimated to control temperature (25 °C), ancestral population acclimated to heat-stress (31 °C), and warm-evolved population acclimated to heat-stress (34 °C). Functional annotation of all identified proteins and gene ontology (GO) enrichment analysis were conducted using the Omicsbox software (version 1.4.12), employing the Fisher exact test with a false discovery rate (FDR) of 0.05. The analysis utilized the reduced option to provide more specific GO terms (Table S4). We compared our simulations against the proteomic dataset qualitatively, based on the up-regulation or down-regulation of specific pathways. This expands our previous work by providing empirical evidence that supports the modeled trade-off between nutrient acquisition and heat-mitigation for both ancestral and evolved populations. The responses of *Prochlorococcus* to warming have been validated previously in Leles and Levine ^23^ against transcriptomic datasets^68^. To validate the model predictions between the two different nutrient regimes, we compared model output to literature values for the proteomic responses of different phytoplankton species between nitrogen-replete and nitrogen-deplete conditions (Table S5, Supplementary Methods). No proteomic datasets were available to validate predictions under combined heat and nutrient stress.

#### Phytoplankton functional types and latitudinal simulations

We solved the proteome model across a latitudinal transect in the Atlantic Ocean, spatially averaging over the longitude range of 330 °to 335 °E, after verifying that temperature and nutrient conditions within this narrow range are relatively uniform. We selected the Atlantic transect for this analysis as it is less dominated by iron limitation compared to the Pacific and therefore offers a more representative environment for examining responses to nitrate limitation. We used historical and future SSP5-8.5 simulations from the CESM2-WACCM model^51^ to construct present-day (1980–2000) and future (2080–2100) averages for temperature and nitrate concentrations, both averaged over the upper 20 m of the water column. Since changes in irradiance were not the focus of this study, we used a constant light dose of 300 *µmol photon m*^−2^ *s*^−1^ as representative of average mixed layer conditions.

We parameterized the model to represent four phytoplankton functional types: *Prochlorococcus*, *Synechococcus*, warm-adapted diatom and cold-adapted diatom. We chose these four types given their importance to global primary production^69^ and their differences in three key properties: 1) nutrient uptake efficiency^45^, 2) thermal traits^50^, and 3) heat-mitigation capacities^46^. Given the wide thermal range of diatoms, we simulated two phenotypes to capture cold-and warm-adapted species. Parameter values were chosen based on previous literature and details are given in the Supplementary Methods (Table S3).

We then expanded our model validation with further experimental data. To confirm that our parameter choices mimic the general physiology of the four types, we validated the emergent maximum growth rates (*µ_max_*) and growth (left) slopes of the thermal curves from our model against data from 167 species previously compiled by Anderson et al. ^50^. Diatom species within this dataset with a *T_opt_* lower than 20 °C were assigned as cold-adapted. All *Prochlorococcus* and *Synechococcus* species in the dataset had a *T_opt_* higher than 20 °C. To make this comparison (Figure 3A), we fitted the data to the Norberg equation. We used a log-likelihood function to fit the parameters of the Norberg equation for each experimental dataset from^50^, then estimated the upper and lower temperature values at which the modeled growth rate was 20 % of its maximum value following^50^. We calculated the lower and upper slopes of the thermal curve by dividing the difference between the maximum growth rate and 20 % of the maximum growth rate value by the difference between the thermally optimum temperature and the identified temperatures at which growth rate was 20 % of maximum.

To parameterize the differential heat-mitigation abilities (*α*) between our functional types, we demonstrate that the modeled concentration of damaged photosystems is approximately 10 times higher for *Prochlorococcus* relative to the warm-adapted diatom, consistent with the findings of Drábková et al. ^46^ (Figure S14). We also performed a sensitivity analysis for the heat-mitigation ability of *Prochlorococcus* for the latitudinal runs (Figure S15), demonstrating that our key finding, i.e. the new potential niche for larger phytoplankton in multi-stressor regions where heat-stress dominates, holds even if we assume equal heat-mitigation abilities among the functional types, but the exact latitudinal range where this happens is variable (Figure S8). Because *Prochlorococcus* has a lower thermal dependency than *Synechococcus* (Table S3), thermal stress develops at lower temperatures when assuming equal heat mitigation abilities. We also compared the emergent averaged cell size and nutrient uptake affinities from our model between cyanobacteria and diatoms to make sure these reflected differences observed empirically (Figure 3B). Finally, we also evaluated if the highest optimal growth rate between cyanobacteria and diatom at any given latitude, as predicted by the proteome model, reflected in higher biomass for that group using simulations from a global model that also considered the different thermal dependencies between these two functional types^25^ (Figure S6).

### QUANTIFICATION AND STATISTICAL ANALYSIS

The computational approach is fully described above and more details can be found in the supplement. Optimization was performed using Ipopt via the JuMP interface in Julia. All data analyses and graphs were performed in R. All code is available in Zenodo.

## SUPPLEMENTAL INFORMATION

Supplemental information can be found online at [add link]

## Supplementary Materials

### Supplementary Methods

#### Model equations

The model is fully defined below. A list of the optimized variables is given in Table S1. Detailed description for each of the metabolic pathways are described in^23^ except for additions/modification to the model which are then described here. Descriptions of the constant parameters are given in the following section as well as Tables S2 and S3.

C1-C8: Steady-state growth of proteins

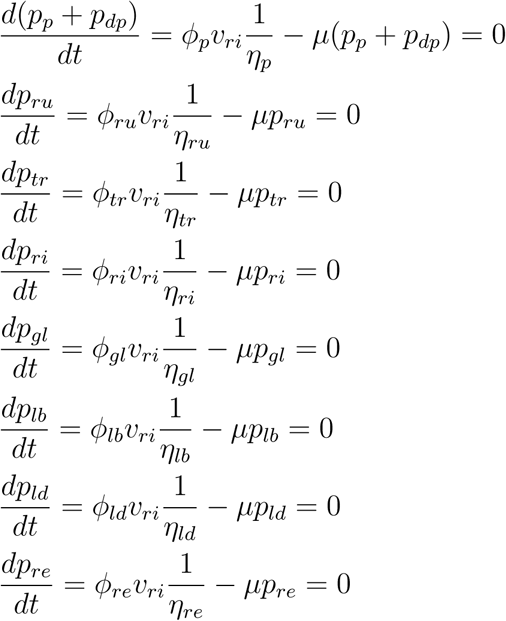

C9-C12: Steady-state growth of other macromolecules

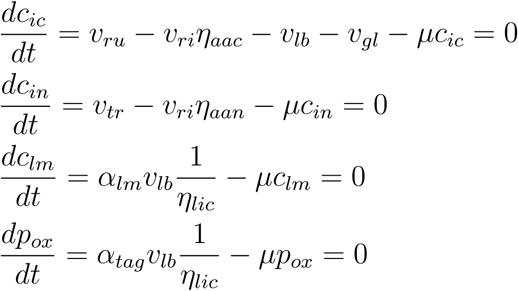

C13: Proteome investments, in which *i* represents protein pools

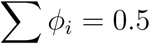

C14: Fractions of the lipid synthesis flux:

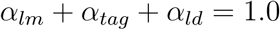

C15: Fraction of damaged photosystems (a function of heat damage and repair)

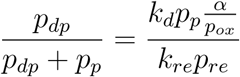

C16: Constraint on the maximum concentration of stored lipids that mitigate damage

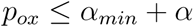

C18: Membrane integrity

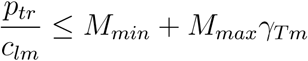

C19: Lipid degradation:

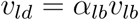

C20-22: Energy metabolism

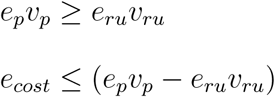

where, *e_cost_* = *e_tr_v_tr_* + *e_ri_v_ri_* + *e_lb_v_lb_* + *e_d_v_d_*+ *e_f_ v_re_*

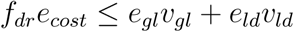

C23: Space constraint

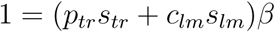

C24: Density constraint, in which *k* indicates all pools of the cell except for photosystems, lipid storage and membrane components

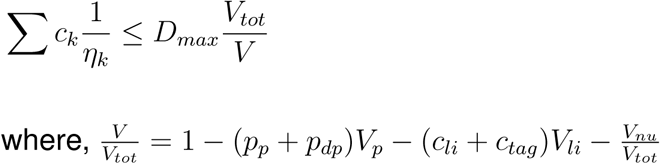

Reaction rates are defined as following:

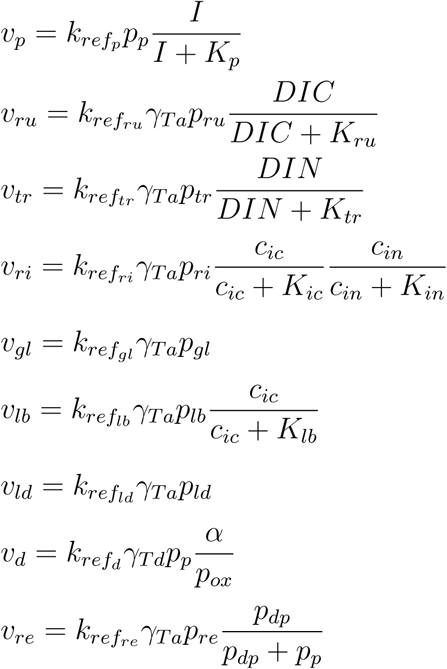

Temperature effects:

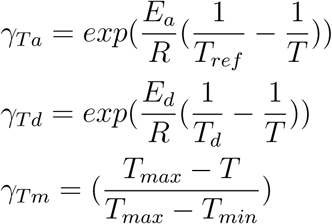

Number of molecules of nitrogen per amino acid:

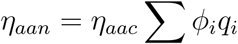

Estimated storage pools that fuel dark respiration:

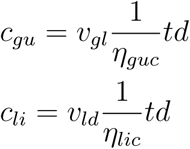

Cell nitrogen to carbon quota (*q_cell_*), in which *i* represents protein pools and *η_i_* represents the molecular weight of proteins in units of aa protein^−1^:

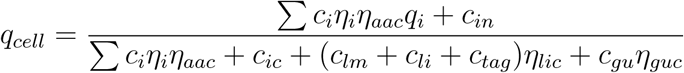

#### Constraining the minimal model cell size

The proteome allocation model solves for the optimal concentration of different macromolecules in the cell given a specific set of environmental conditions. We impose a constraint on the maximum intracellular density of macromolecules (*D_max_*) such that, if the optimal metabolic strategy involves building more proteins and the cell is at *D_max_*, the cell must increase its cell size. The volume of the cell is defined by the volume occupied by photosystems and stored lipids as well as all other macromolecules, following Equations 41-43 in^23^. These constraints effectively keep cell size from increasing indefinitely in our model. In order to constraint the minimal size of our modeled cell, we require the model to account for the intracellular space taken up by the cellular nucleus (*V_nu_*). Thus we modify Equation 41 from^23^:

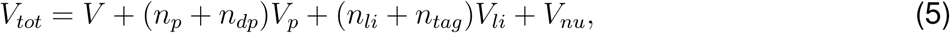

in which *V_tot_* is the total cellular volume, *V* is the volume occupied by all macromolecules with the exception of photosystems, stored lipids, and nucleus. *n_p_* and *n_dp_* are the total number of functional and damaged photosystems, respectively, and *n_li_* and *n_tag_* are the total number of stored lipids required to regulate oxidative stress and to fuel respiration, respectively. *V_p_* and *V_li_* correspond to the volume of photosystem and stored lipid molecules, respectively.

#### Parameter values

The parameters used to define the different phytoplankton functional types can be found in Table S3. Below we describe in detail our choices for each of these parameters. All other parameter values of the model were kept the same and can be found in Table S2; for detailed description on the choice of these parameter values and respective references please refer to the supplemental methods in Leles and Levine ^23^.

*E_a_*: this parameter sets the slope at which metabolic rates in the model increase exponentially as a function of temperature. To parameterize *E_a_*, we derived the growth (left) slope of the thermal growth curves for the different functional types from empirical data available for 167 species of phytoplankton based on the compilation in^50^ and validated our model against the averaged values obtained from empirical observations (Figure 3A).

*T_d_*: similarly, the heat-stress temperature was defined based on averaged values from the same dataset compiled in Anderson et al. ^50^.

*K_tr_*: the half-saturation for nitrogen uptake was derived from Figure 1B in Edwards et al. ^45^.

*V_p_*: the average volume occupied by a photosystem molecule is 3 *×* 10^−5^*µm*^3^ for *Thalassiosira pseudonana* ^70^ and between 1 *×* 10^−6^ - 2 *×* 10^−5^ *µm*^3^ for the cyanobacteria *Synechococcus* (BioNumbers: ID103908 and ID103909).

*V_nu_*: the nucleus volume was derived from^71^, in which the nucleus volume was given for several species of phytoplankton.

*α*: the maximum concentration of antioxidants (i.e. stored lipids) in the cell (not used to fuel dark respiration) was constrained based on the average fraction of the cellular volume that is occupied by lipids, i.e. 20%^72^, resulting in a maximum value of 2 × 10^7^ molecules *µm*^−3^.

*e_d_*: we assumed that the rate of heat-damage scales with 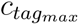 so that the cost is higher for cyanobacteria based on their higher sensitivity to heat stress^46^, as following: 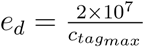.

#### Settings for the sensitivity analyses

The baseline parameters used for the sensitivity analyses (Figures 2D-G) were the same as those set for the warm-adapted diatom type, with the exception of the following parameters:

*K_tr_*: 10.0 *µM*
*E_a_*: variable, 10 values from 0.55 to 1.05 eV
*α*: variable, 10 values from 6 *×* 10^6^ to 2.4 *×* 10^7^ molecules *µm*^−3^.

We changed the value of *K_tr_* to represent cells that have higher nutrient affinity than our modeled diatoms, but lower nutrient affinity than our cyanobacteria types (Table S3). By varying *E_a_* and *α* we were able to capture phenotypes with diverse strategies, from varying degrees of thermal dependency, as well as cells with different abilities to mitigate oxidative stress caused by heat-damage.

The phenotypes shown in Figure 2A assumed a fixed *α* of 2 × 10^7^ and variable *E_a_* while the phenotypes shown in Figure 2B assumed a fixed *E_a_* of 0.55 and variable *α*. The two example phenotypes shown in Figure 2C differed in the following parameters:

Nutrient-efficient: *E_a_* = 1.05 and *α* = 6 *×* 10^6^
Thermal-tolerant: *E_a_*= 0.61 and *α* = 2.4 *×* 10^7^.

The environmental conditions for these analyses were set as following:

Temperature: from 18 to 35 °C, every 0.25 °C

Dissolved Inorganic Nitrogen (DIN) concentrations: replete (20 *µM*) and limited (0.1 *µM*) conditions

Irradiance: 300 *µmol photon m*^−2^ *s*^−1^.

#### Improving optimization algorithm

The proteome model solves for the optimal steady-state solution which maximizes growth rates. However, sometimes the optimizer can get stuck in a local minimum instead of finding the global minimum. This can occur for many reasons including the optimizer ‘timing out’ due to the optimization surface being too flat or poor initial conditions. As this work required solving for 10,000 combinations of trait values (model parameters) and environmental conditions, we needed to develop a pipeline for ensuring that the optimizer indeed found the global minimum. We used an iterative approach based on the assumption that the model solutions would be either continuous or exhibit a relatively small number of states across the tested environmental conditions.

We first conducted a set of simulations across the environmental gradient (e.g., temperature) using several random initial guesses. We used the optimized solutions from these initial runs to identify ‘anomalous’ solutions that might be local minima. Specifically, we used a moving average approach relying on tangent line calculations to evaluate the rate of change between consecutive solutions across a temperature gradient. An abrupt change in the tangent between consecutive data points resulted in the solution being flagged as potentially ‘anomalous’. Solutions which violated biologically realistic bounds for key variables were also flagged. We used the following bounds for each of the key variables: growth rates (0.001-2 *day*^−^1), cell length (0.1-30 *µm*), and nitrogen to carbon quota of the cell (0.008-0.5 mol N:mol C). We then identified, for each ‘anomalous’ solution, the closest preceding and succeeding ‘good’ solution (i.e., solutions not flagged as anomalous). The initial conditions (starting guesses) for the optimizer were then derived by extrapolating between these closest good solutions using a linear interpolation. We performed two re-optimization passes using this strategy. For each pass, the new solutions were compared against the original ‘anomalous’ solution and the one with the maximum growth rate was selected (the optimizer cost function maximizes growth rates). This allows us to reduce the impact of local minimum and ensure the results presented were indeed the optimal (global) solution.

#### Model comparison against proteomic data

We compared the proteomic shifts predicted by the model against proteomic data previously published in the literature for diatom and cyanobacteria species grown under controlled growth conditions. We also compared the shifts in response to warming predicted by the model to data from an experimental evolution study^73^. This dataset provided robust evidence supporting the trade-off between investing in transporters versus glycolysis and heat-mitigation strategies (Table S4). The GO terms used for this comparison are given in Table S4. We were unable to find any previous studies that reported the proteomic response of phytoplankton to the multi-stressor growth condition of warming and nutrient limitation. Therefore, we compared the model against other datasets to evaluate the metabolic responses to only nutrient limitation (Table S5). The list of proteins/gene terms used for this comparison is specified below:

Nutrient trasporters: nitrate/ammonia transporters^74^, nitrate/ammonium assimilation^74^, phosph binding protein^75^, nitrogen assimilation enzymes (nitrate reductase and glutamate synthase)^28^, nitrate/nitrite transporters^76^, and phosphate transporters^76^.

Photosystems: proteins from^75^ are described in their Table 1 and correspond to Cytochrome b559 subunit beta, Photosystem II reaction center protein L, Cytochrome b6-f complex subunit 4, Cytochrome b559 subunit alpha, Divinyl chlorophyll a/b light-harvesting protein PcbE, Divinyl chlorophyll a/b light-harvesting protein PcbD. Others included Fucoxanthin chlorophyll a/c protein^28^. See also Figure 5B for detailed proteins in^76^ and Figure 5 and Supplementary file I in^74^.

Rubisco: Ribulose bisphosphate carboxylase and other proteins related to carbon fixation; see Figure 5B for detailed proteins in^76^ and Figure 5 and Supplementary file II in^74^.

Ribosomes: ribosomal proteins (small and large units); see Figure 5D in^76^.

Glycolysis: KEGG pathways related to Glycolysis/Gluconeogenesis, which includes Embden-Meyerhof pathway (glucose to pyruvate), glycolysis, pyruvate oxidation and gluconeogenesis; see Table S5 in^76^.

Lipid synthesis: KEGG pathways associated with Glycerophospholipid metabolism (see Table S5 in^76^), and Glycerolipid metabolism, Fatty acid metabolism and elongation (see Supplementary file II in^74^).

### Supplemental Figures

**Figure S1:**
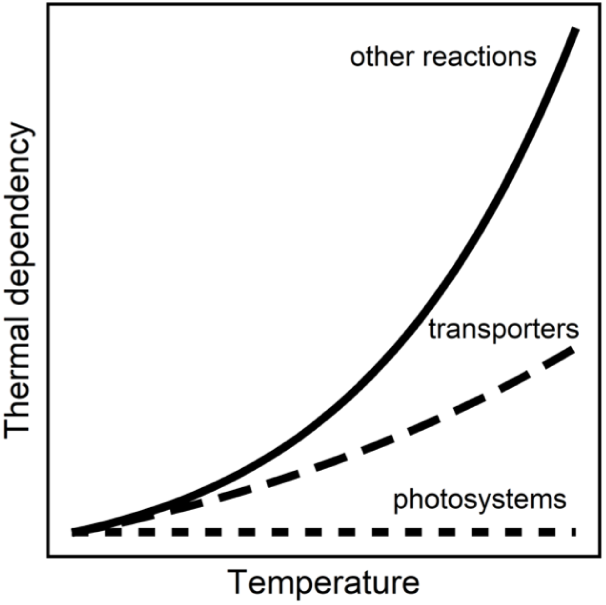
Differential temperature sensitivity of metabolic reactions captured in our proteome allocation model. Light-dependent reactions driven by photosystems are assumed to be temperature-independent. Nitrogen transporters are assumed to have a lower thermal sensitivity compared to other reactions in the model, such as respiration (mediated by glycolysis proteins), protein synthesis (mediated by ribosomes), and carbon fixation (mediated by the rubisco pool).

**Figure S2:**
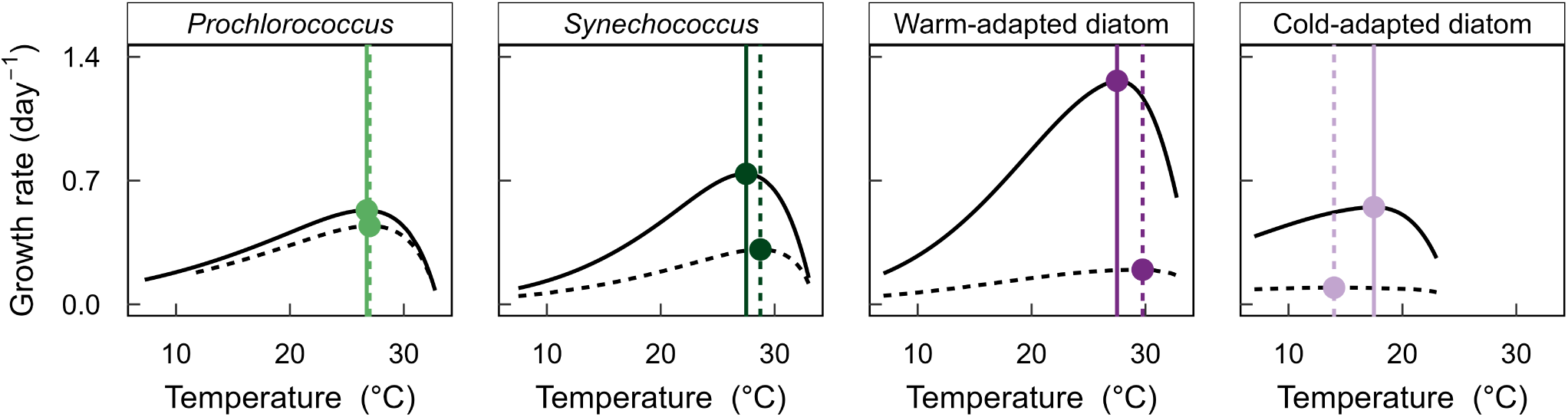
Emergent thermal growth curves for the different phytoplankton functional types under nitrogen-replete (solid lines; 20 *µ*M) and nitrogen-limiting (dashed lines; 0.1 *µ*M) conditions. Symbols and vertical lines indicate temperature optima (*T_opt_*) in both nutrient regimes. Colors indicate the different phytoplankton functional types following Figures 3 and 4.

**Figure S3:**
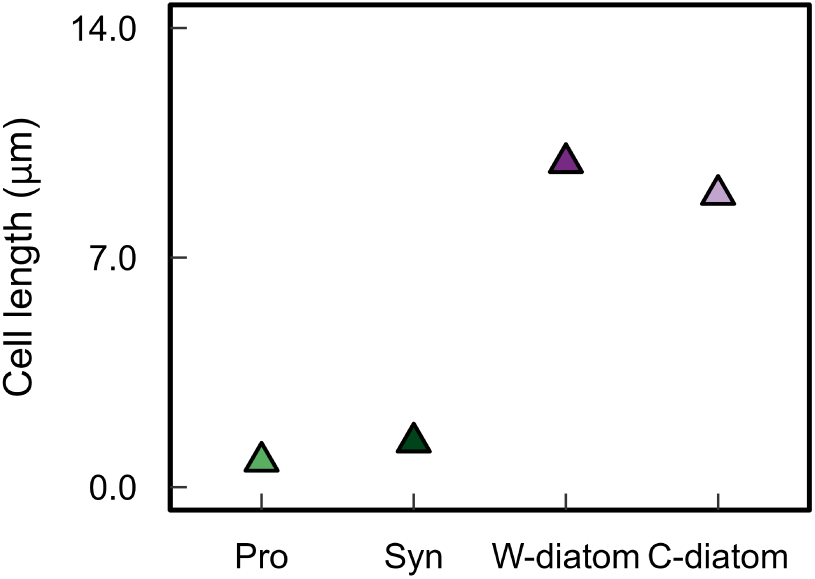
Cell diameter under nitrogen replete (20 *µ*M) conditions. Data are given at *T_opt_* for each functional type as following: Pro: *Prochlorococcus*, Syn: *Synechococcus*, W-diatom: warm-adapted diatom, C-diatom: cold-adapted diatom. Colors indicate the different phytoplankton functional types following Figures 3 and 4.

**Figure S4:**
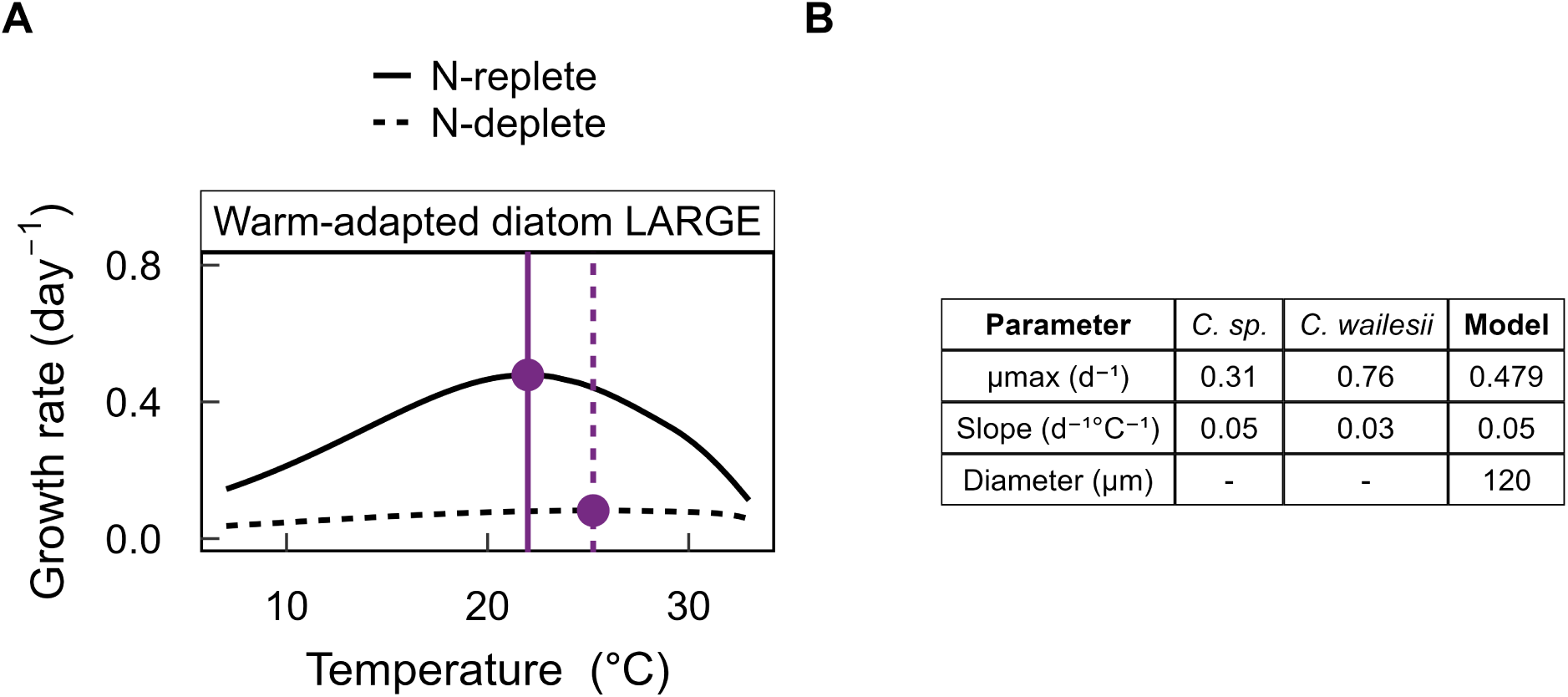
Simulated large warm-adapted diatom type. The only differences in parameterization from the warm-adapted diatom type were: *V_nu_* = 2.5 × 10^4^ *µm*^3^ and *D_max_* = 1.8 × 10^9^ Da *µm*^−3^. A) Modeled thermal growth curve for N-replete (20 *µ*M) and N-limited (0.1 *µ*M) conditions. Symbols and vertical lines indicate temperature optima (*T_opt_*) in both nutrient regimes. B) Model validation against empirical data for two species of large diatoms (*>* 100 *µ*m) of the genus *Coscinodiscus*. Empirical data was obtained from Anderson et al. ^50^.

**Figure S5:**
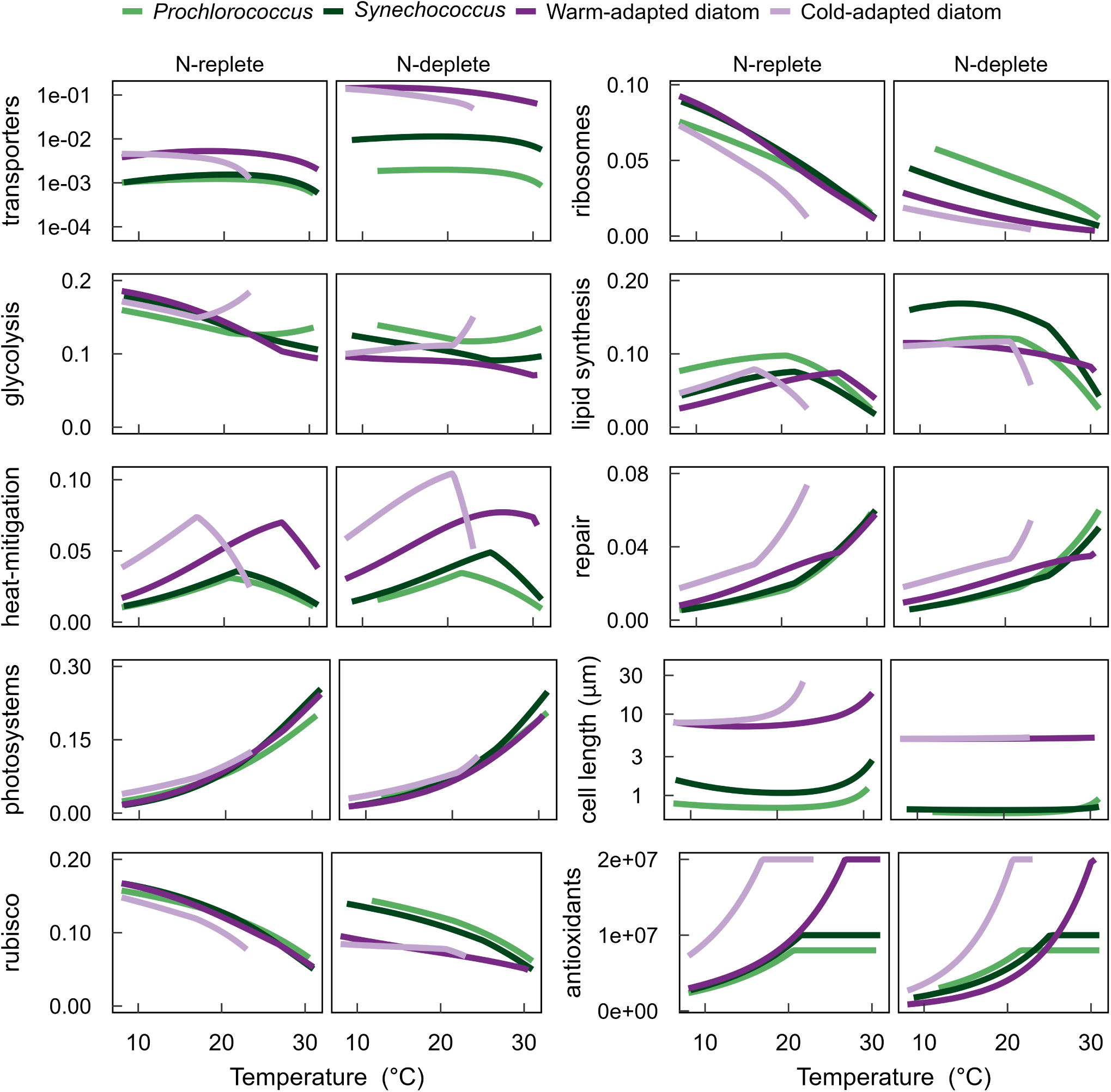
Optimal relative proteome investments and cell size. Model output as a function of temperature for N-replete (20 *µ*M) and N-limited (0.1 *µ*M) conditions are compared among four phenotypes corresponding to those illustrated in Figures 3 and 4. Relative proteome investments vary between 0 and 1 and antioxidants are given in units of number of molecules *µm*^−3^.

**Figure S6:**
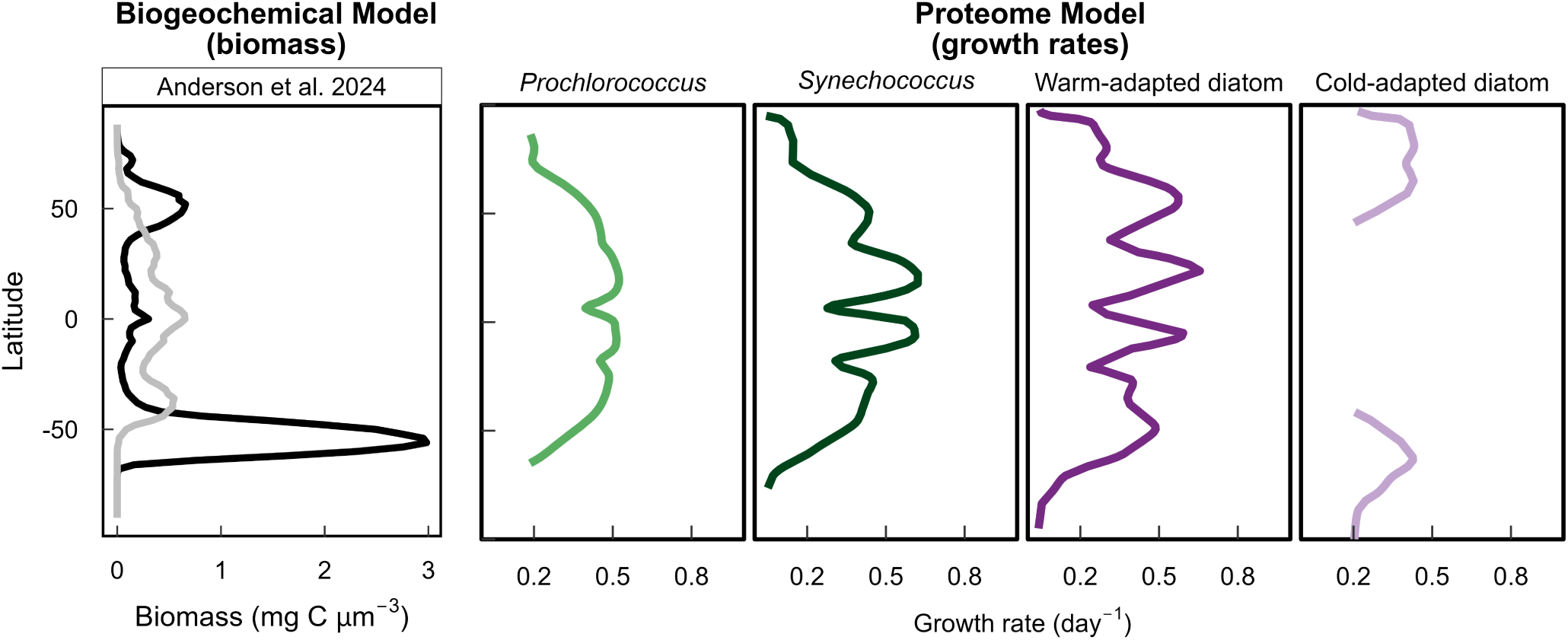
Mean phytoplankton biomass across a latitudinal gradient from the DARWIN model. Integrated biomass over the upper 240 m averaged over 10 years is given for diatoms (black line) and cyanobacteria (gray line). Simulated data were obtained from^25^. Also shown are the optimal growth rates from the proteome model for the different functional types.

**Figure S7:**
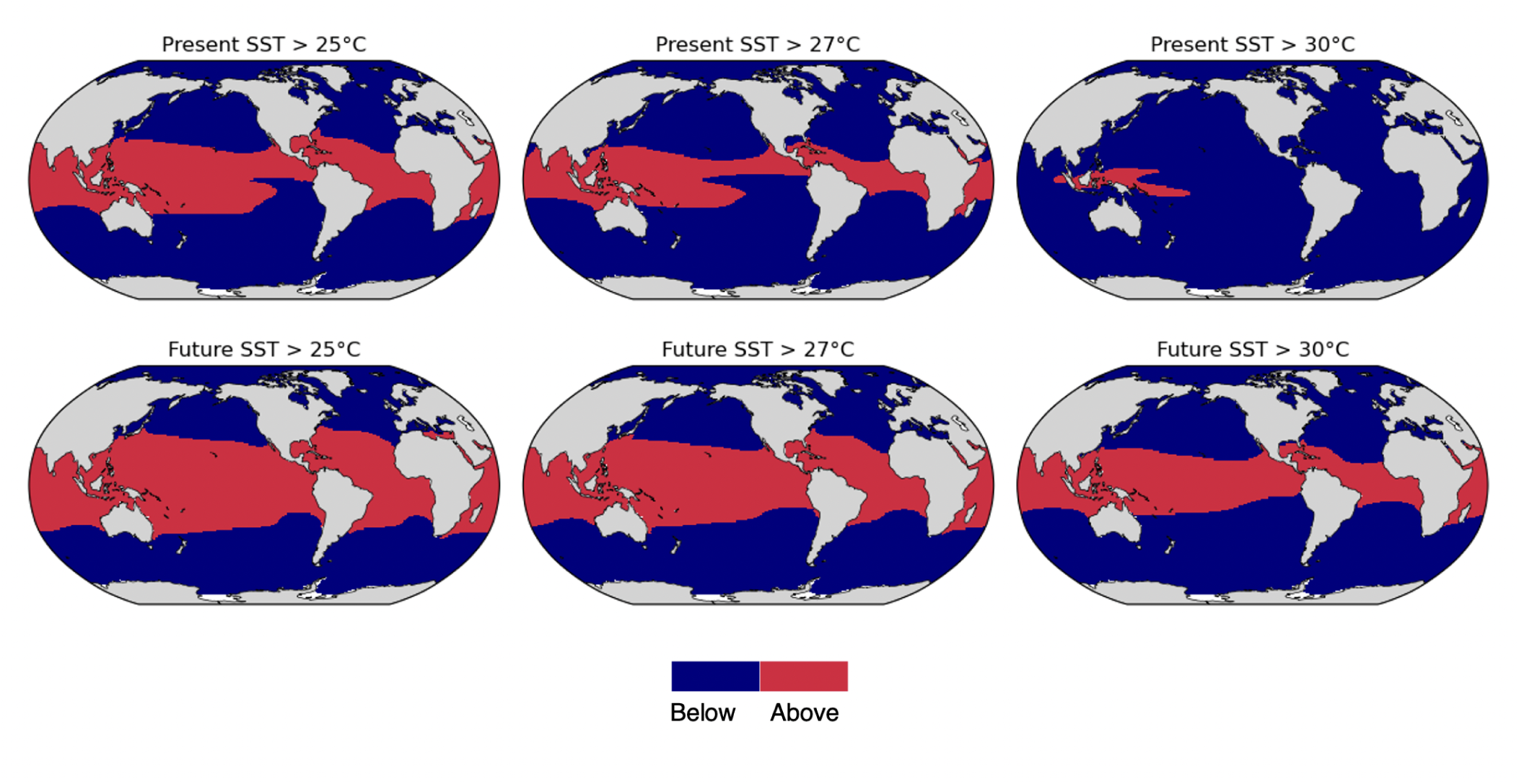
Projections of oceanic regions experiencing different surface sea temperature (SST) thresholds under present-day and future climate scenarios from the CESM2-WACCM model under SSP5-8.5. Under the future climate scenario, the model predicts a global increase in surface waters experiencing temperatures above: 25°C from 37.7% to 52.9%, 27°C from 26.9% to 46.3%, and 30°C from 2.7% under present day to 34.5% by the end of the century. Temperature thresholds were chosen based on the *T_opt_* of the different modeled phytoplankton functional types (Figure S2).

**Figure S8:**
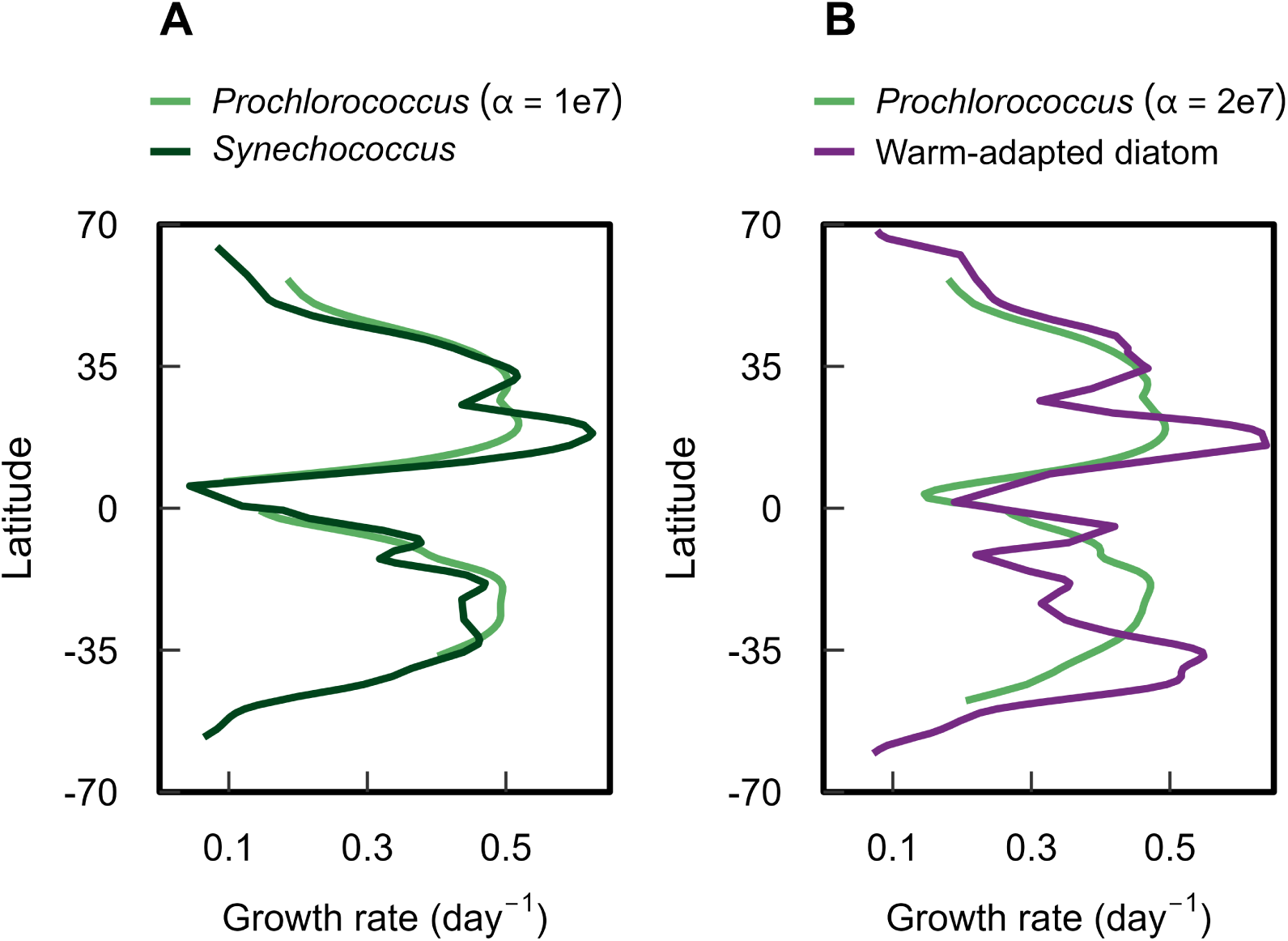
Sensitivity analysis to the heat-mitigation ability of *Prochlorococcus*. Our baseline simulation assumes that *Prochlorococcus* has lower heat-mitigation ability than the other functional types, with *α* = 8*e*6 (Figure 4 and Table S3). Here we assume all else equal and only vary *α* so that it is the same as *Synechococcus* (A) or the warm-adapted diatom (B).

**Figure S9:**
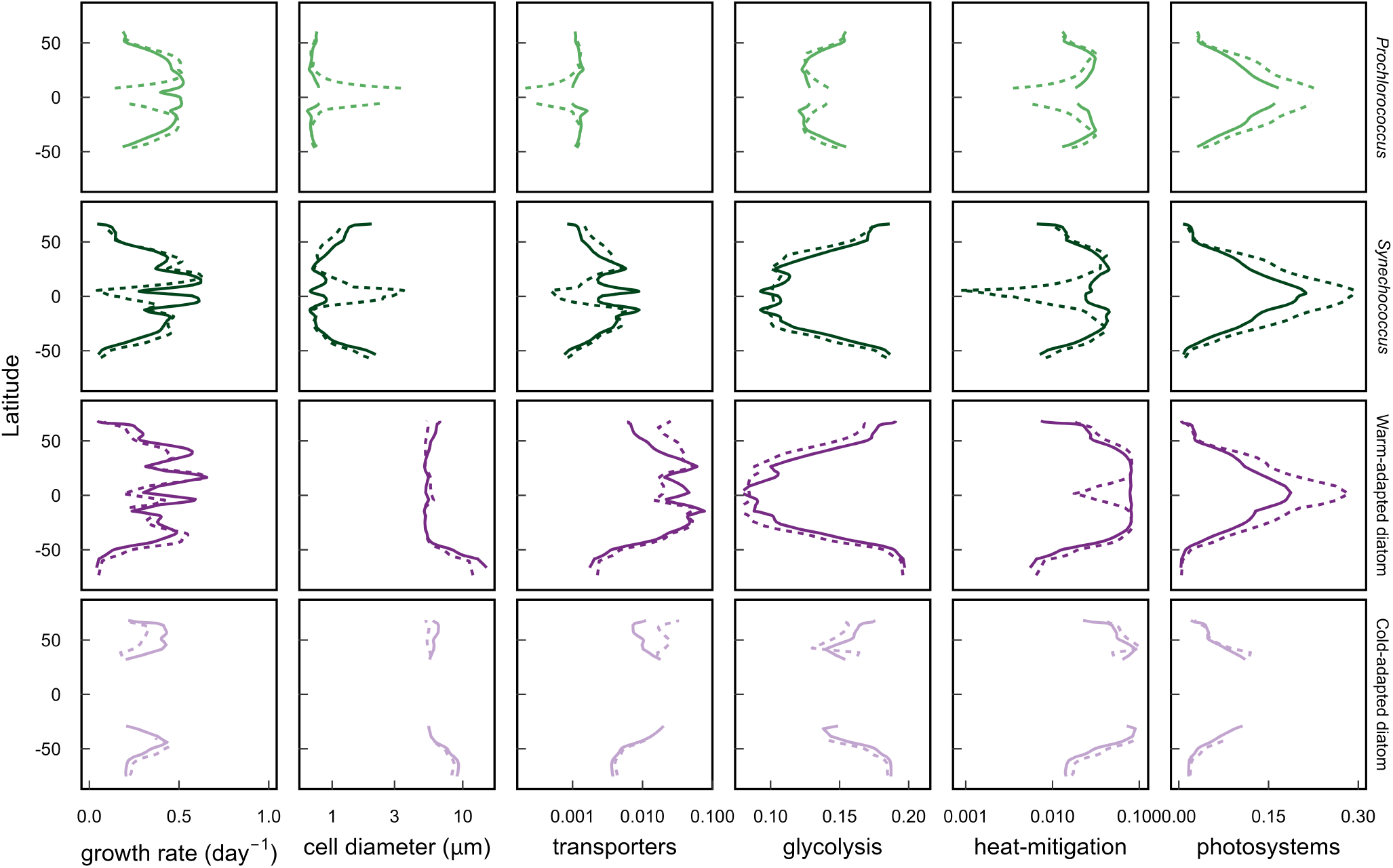
Phytoplankton optimal metabolic strategies across a latitudinal transect in the Atlantic Ocean based on temperature and nitrogen concentrations averaged over the upper 20 m of the water column. Simulations were performed for present-day (1980-2000; solid lines) and future (2080-2100; dashed lines) projections of nitrate concentrations and temperature using model output from the the CESM2-WACCM model under SSP5-8.5 (Figure 4B). Results are given for four phytoplankton functional types for optimal growth rates, cell diameter, and relative proteome investments that vary between 0 and 1. Note that no data is shown when growth rates were equal to zero.

**Figure S10:**
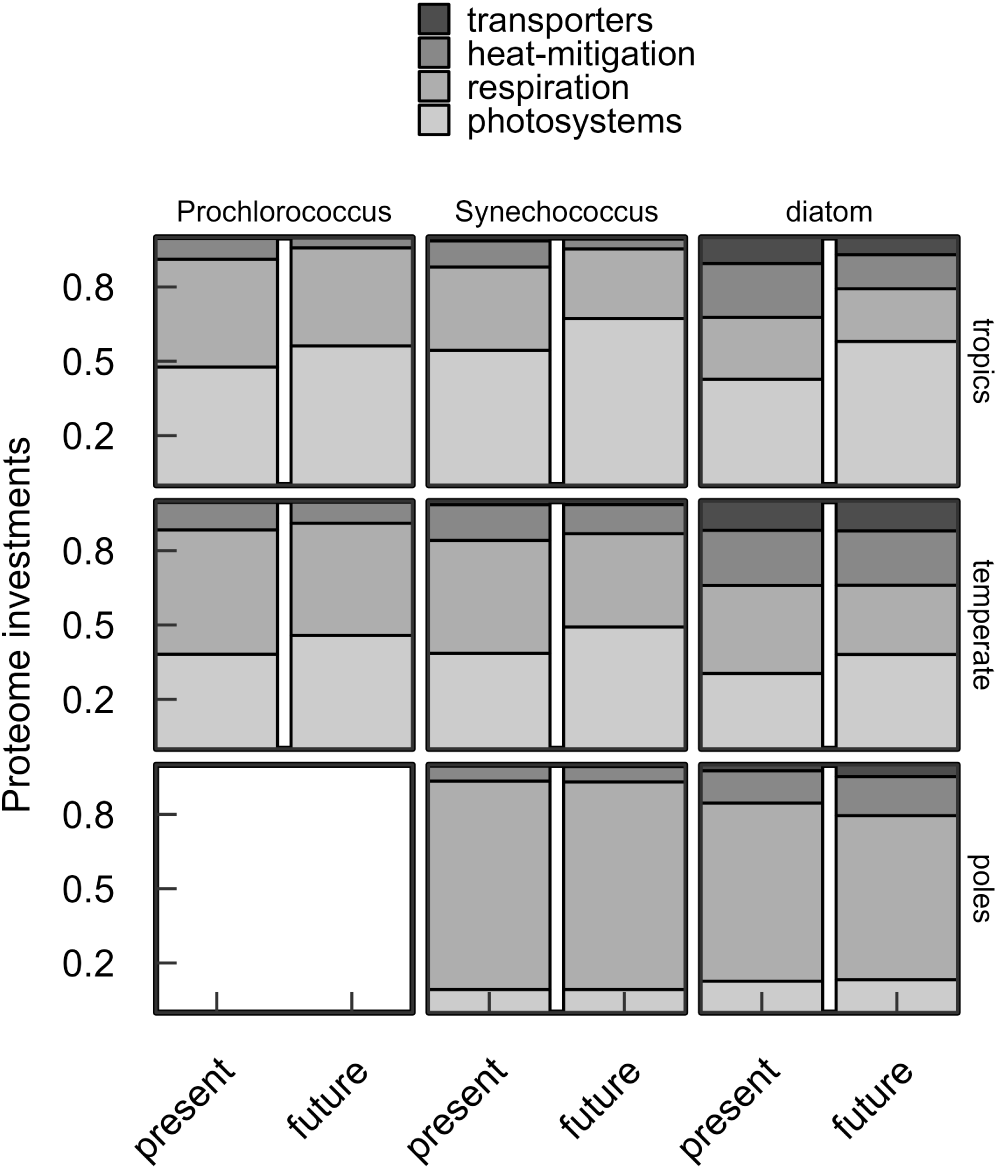
Metabolic strategies based on phytoplankton functional type across major oceanic regions. Scaled relative proteome investments (varies between 0 and 1) for present-day and future climate projections across three distinct regions of the ocean: tropics (22°S-22°N), temperate (25°-45°) and poles (50°-70°). For full latitudinal simulations, see Figure S9.

**Figure S11:**
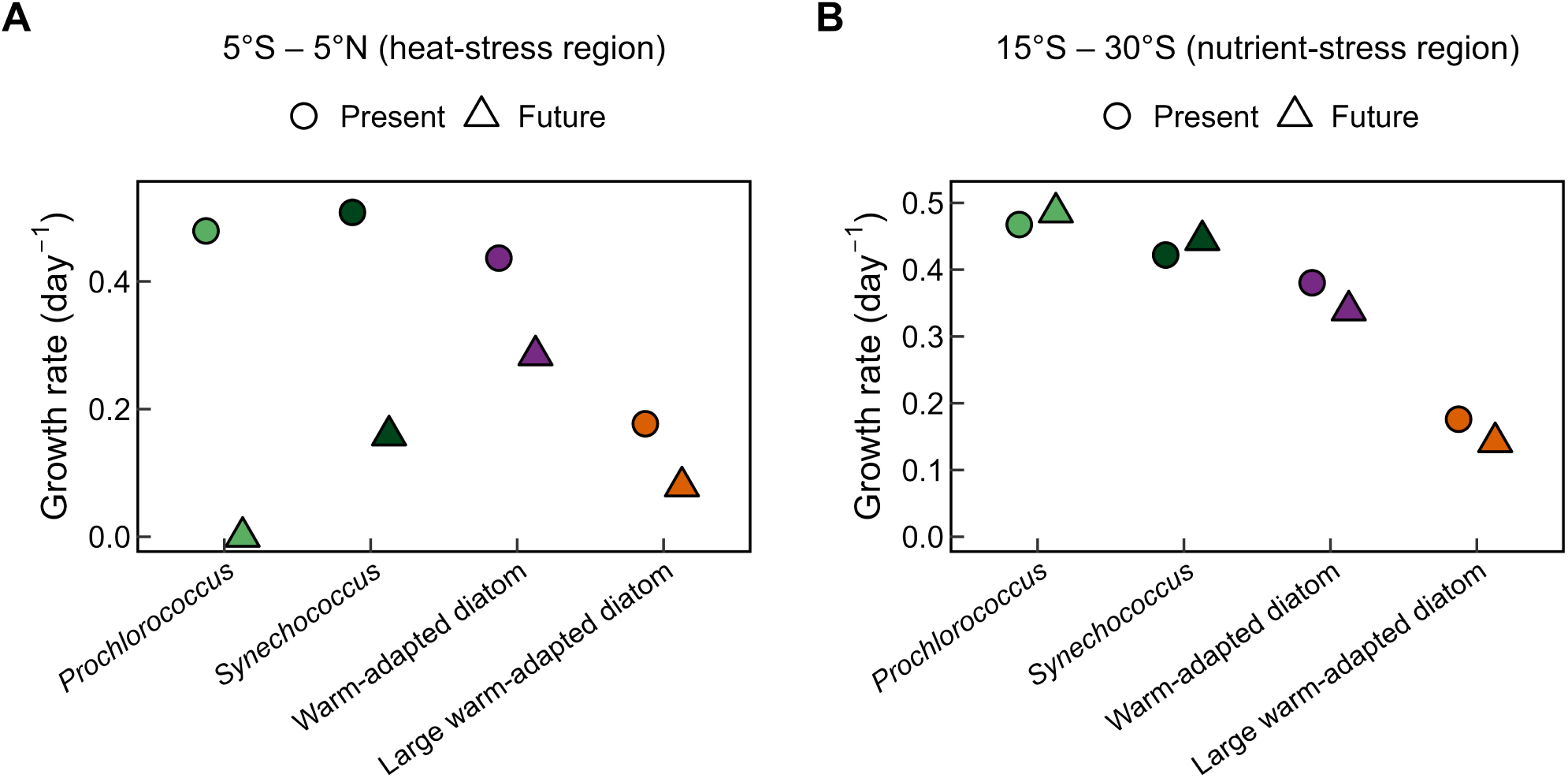
Averaged growth rates under multi-stressor growth regions. The latitudinal ranges 5°S-5°N and 15°S-30°S were selected as examples of ‘heat-stress’ and ‘nutrient-stress’ regions, respectively.

**Figure S12:**
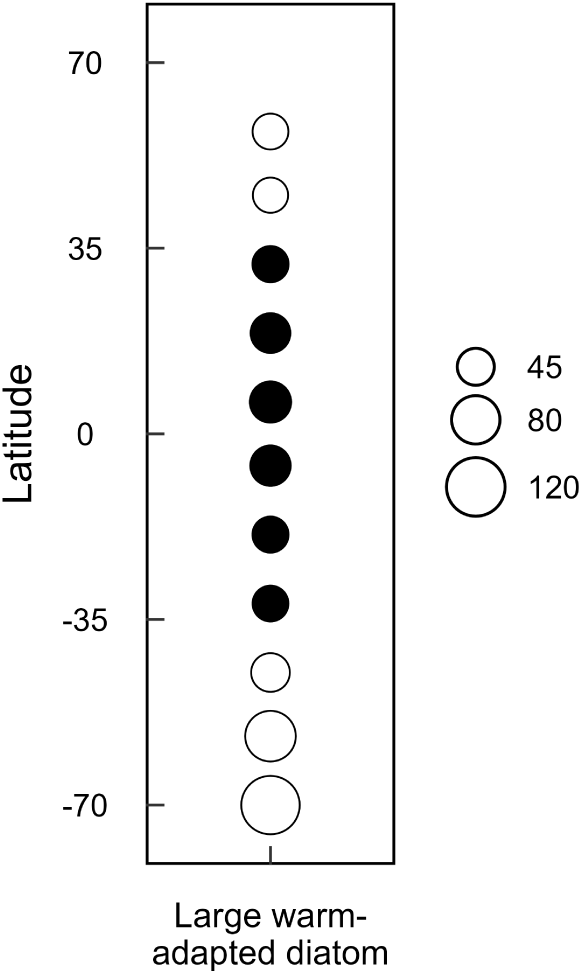
Optimal cell diameter (*µm*) under the future climate scenario for the large warm-adapted diatom. White fill and black fill shows decreases and increases in cell size from present to future, respectively.

**Figure S13:**
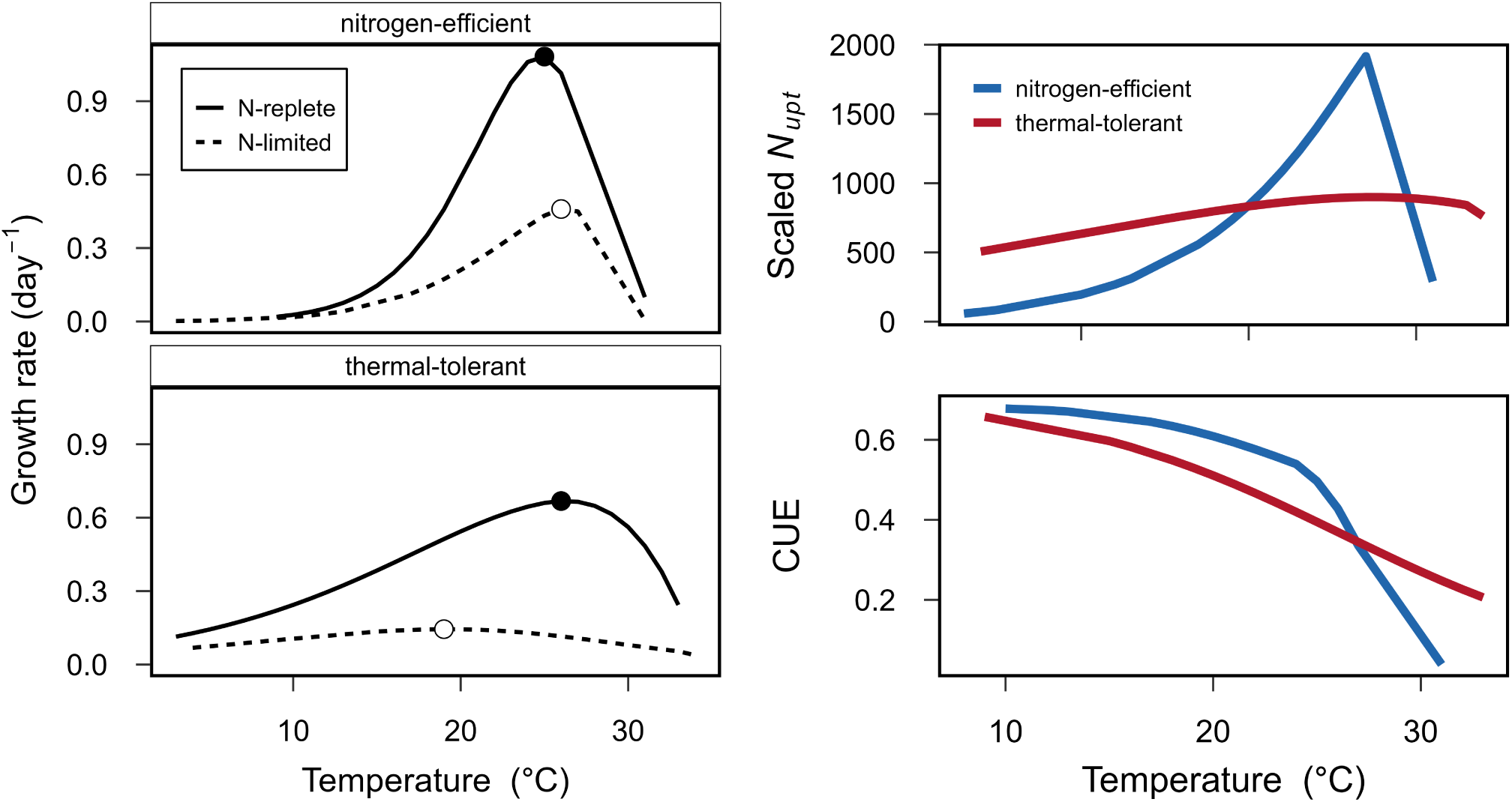
Model runs where phytoplankton produce antioxidant enzymes to decrease oxidative stress instead of lipid storage in the form of TAGs. (A) thermal growth curves indicate shifts in *T_opt_* from nitrogen-replete to nitrogen-limited conditions as a result of different metabolic strategies. (B) model nitrogen uptake rates (*N_upt_*) and Carbon Use Efficiencies (CUE) under nitrogen limitation. Carbon use efficiency is defined as 1 - respiration/photosynthesis.

**Figure S14:**
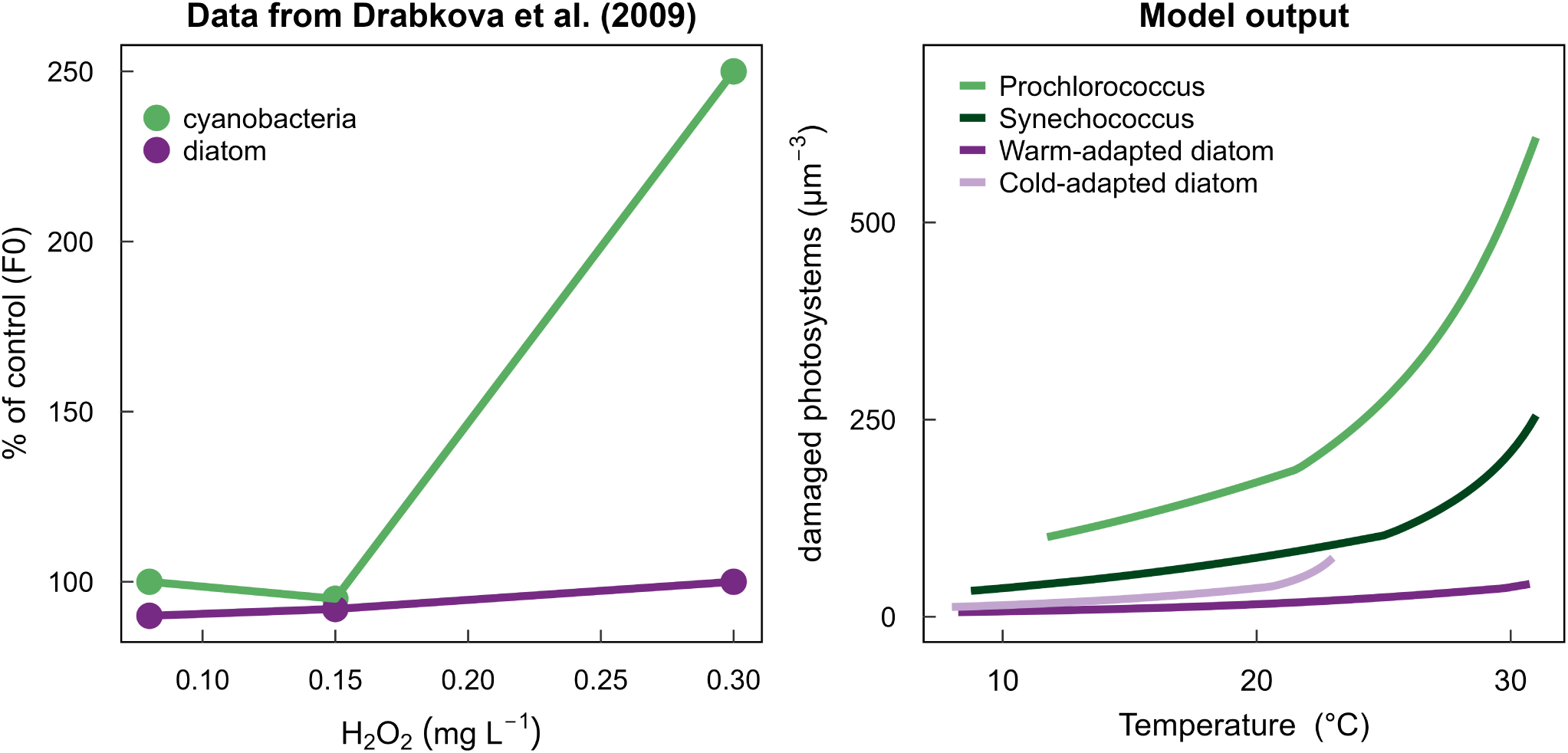
Qualitative validation of sensitivity to oxidative stress among different functional types. Left panel shows F_0_ as a percentage of control for cultures of cyanobacteria and diatoms under increasing H_2_O_2_ concentrations, redrawn from Drábková et al. ^46^. Right panel shows modeled concentration of damaged photosystems as a function of temperature under N-deplete conditions. Consistent with the empirical observations, the concentration of damaged photosystems are substantially higher in *Prochlorococcus* than in the warm-adapted diatom across the full temperature range, capturing the 10-fold differential ROS sensitivity reported by Drábková et al. ^46^.

**Figure S15:**
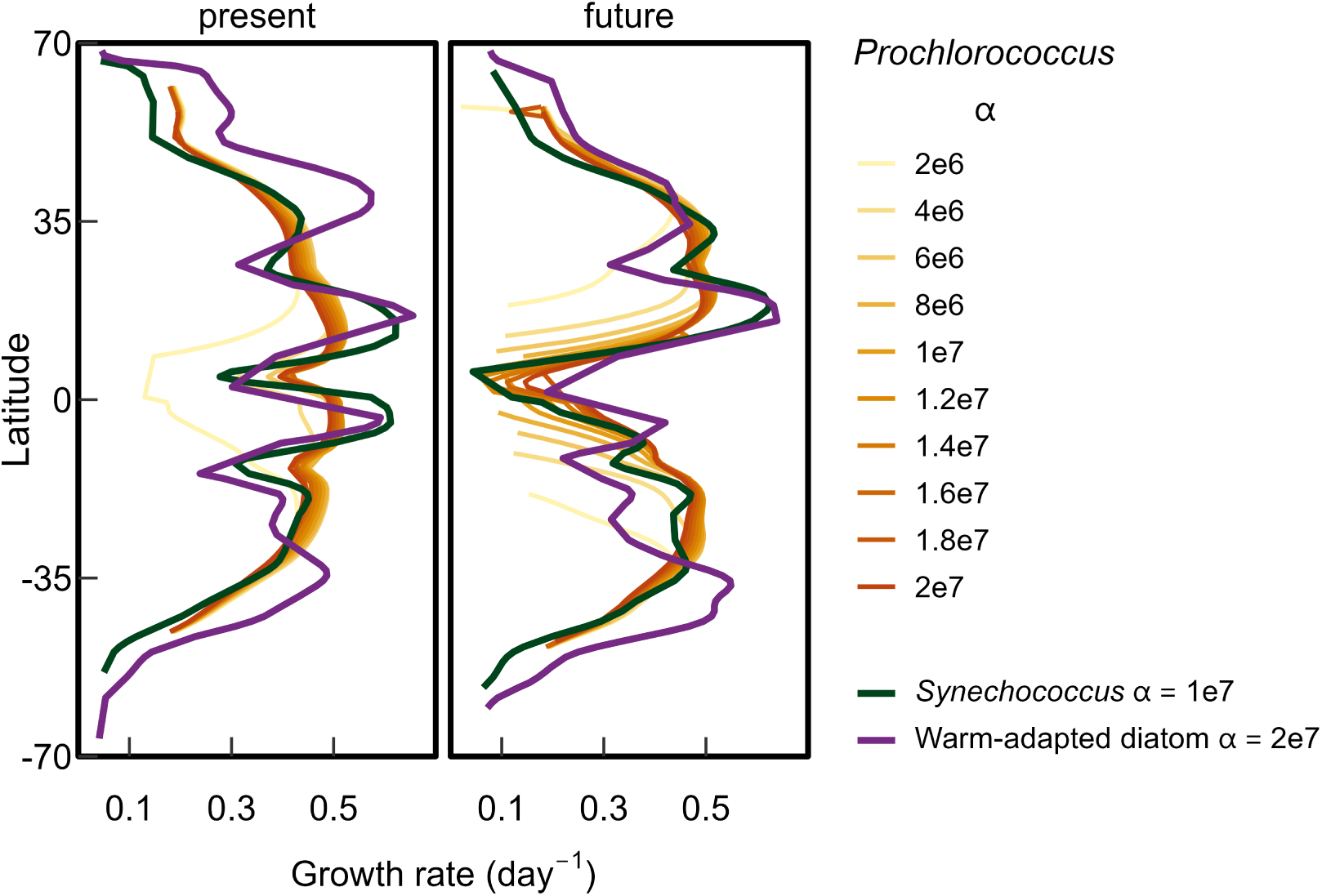
Sensitivity analysis to the heat-mitigation ability of *Prochlorococcus* across a range of values. Our baseline simulation assumes *α* = 8*e*6 for *Prochlorococcus* (Figure 4 and Table S3). Overlaid are the potential growth rates for our baseline runs for *Synechococcus* and the warm-adapted diatom for comparison.

### Supplemental Tables

**Table S1:** Optimized parameters. Abbreviations, descriptions and units for the optimized variables. State variables correspond to the concentration of proteins and other macromolecules inside a phytoplankton cell.

| State Variables |  |  |
| --- | --- | --- |
| Proteins |  |  |
| $p_p$ | Photosystems | molecules of $p_p \mu m^{-3}$ |
| $p_{ru}$ | rubisco | molecules of $p_{ru} \mu m^{-3}$ |
| $p_{tr}$ | Transporters | molecules of $p_{tr} \mu m^{-3}$ |
| $p_{ri}$ | Ribosomes | molecules of $p_{ri} \mu m^{-3}$ |
| $p_{gl}$ | Glycolysis | molecules of $p_{gl} \mu m^{-3}$ |
| $p_{lb}$ | Lipid Synthesis Pathway | molecules of $p_{lb} \mu m^{-3}$ |
| $p_{ld}$ | Lipid Degradation Pathway | molecules of $p_{ld} \mu m^{-3}$ |
| $p_{re}$ | Repair Proteins | molecules of $p_{re} \mu m^{-3}$ |
| Other macromolecules |  |  |
| $c_{ic}$ | Internal Carbon | molecules of C $\mu m^{-3}$ |
| $c_{in}$ | Internal Nitrogen | molecules of N $\mu m^{-3}$ |
| $c_{lm}$ | Membrane Lipids | molecules of $c_{lm} \mu m^{-3}$ |
| $p_{ox}$ | Antioxidants (as lipid storage) | molecules of $p_{ox} \mu m^{-3}$ |
| Parameters |  |  |
| Relative proteome investments in proteins |  |  |
| $\phi_p$ | Photosystems | unitless |
| $\phi_{ru}$ | rubisco | unitless |
| $\phi_{tr}$ | Transporters | unitless |
| $\phi_{ri}$ | Ribosomes | unitless |
| $\phi_{gl}$ | Glycolysis | unitless |
| $\phi_{lb}$ | Lipid Synthesis Pathway | unitless |
| $\phi_{ld}$ | Lipid Degradation Pathway | unitless |
| $\phi_{re}$ | Repair Proteins | unitless |
| Other parameters |  |  |
| $\beta$ | Volume to Surface Area ratio of the cell | $\mu m$ |
| $\alpha_{lm}$ | Fraction of lipid synthesis flux used to build membrane lipids | unitless |
| $\alpha_{ld}$ | Fraction of lipid synthesis flux used to fuel lipid degradation | unitless |
| $\alpha_{tag}$ | Fraction of lipid synthesis flux used to build stored lipids | unitless |

**Table S2:** Parameter values. Abbreviations, descriptions, values, and units. The values of tunable parameters are given in Table S3. All other parameter values were kept constant across our simulations. See Supplementary Text in^23^ for references and more details regarding the source of the parameter values.

| Maximum reference turnover rates (# molecules per protein per time) |  |  |  |
| --- | --- | --- | --- |
| $k_{ref_p}$ | Photosystems | 7800 | photons $\text{min}^{-1}$ |
| $k_{ref_{ru}}$ | rubisco | 157 | molecules of C $\text{min}^{-1}$ |
| $k_{ref_{tr}}$ | Transporters | 120 | molecules of N $\text{min}^{-1}$ |
| $k_{ref_{ri}}$ | Ribosomes | 114 | molecules of aa $\text{min}^{-1}$ |
| $k_{ref_{gl}}$ | Glycolysis | 120 | molecules of C $\text{min}^{-1}$ |
| $k_{ref_{lb}}$ | Lipid Synthesis | 120 | molecules of C $\text{min}^{-1}$ |
| $k_{ref_{ld}}$ | Lipid Degradation | 120 | molecules of C $\text{min}^{-1}$ |
| $k_{ref_d}$ | Heat damage | 1000 | molecules of $p$ $\text{min}^{-1}$ |
| $k_{ref_{re}}$ | Repair Proteins | 1000 | molecules of $p$ $\text{min}^{-1}$ |
| Half saturation constants |  |  |  |
| $K_p$ | Photosystems | 10 | $\mu\text{mol photon } m^{-2} s^{-1}$ |
| $K_{ru}$ | rubisco | 1 | $\mu\text{M DIC}$ |
| $K_{tr}$ | Transporters | Table S3 | $\mu\text{M DIN}$ |
| $K_{in}$ | Nitrogen Metabolism | 10000 | molecules of N $\mu m^{-3}$ |
| $K_{ic}$ | Carbon Metabolism | 10000 | molecules of C $\mu m^{-3}$ |
| $K_{gl}$ | Glycolysis | 10000 | molecules of C $\mu m^{-3}$ |
| $K_{lb}$ | Lipid Synthesis | 10000 | molecules of C $\mu m^{-3}$ |
| $K_{ld}$ | Lipid Degradation | 10000 | molecules of C $\mu m^{-3}$ |
| Temperature effects |  |  |  |
| $E_a$ | Thermal dependency | Table S3 | eV |
| $E_{atr}$ | $E_a$ of transporters | $E_a/2$ | eV |
| $E_d$ | Deactivation energy | 1.4 | eV |
| $T_d$ | Stress temperature | Table S3 | $^{\circ}\text{C}$ |
| $T_{ref}$ | Reference temperature | 20 | $^{\circ}\text{C}$ |
| $R$ | Boltzmann's constant | $8.62 \times 10^{-5}$ | eV/K |
| Molecular weights |  |  |  |
| $\eta_p$ | Photosystems | 9082 | aa/molecule of $p$ |
| $\eta_{ru}$ | rubisco | 5933 | aa/molecule of $ru$ |
| $\eta_{tr}$ | Transporters | 1046 | aa/molecule of $tr$ |
| $\eta_{ri}$ | Ribosomes | 19393 | aa/molecule of $ri$ |
| $\eta_{gl}$ | Glycolysis | 15220 | aa/molecule of $gl$ |
| $\eta_{lb}$ | Lipid Synthesis | 13742 | aa/molecule of $lb$ |
| $\eta_{ld}$ | Lipid Degradation | 5128 | aa/molecule of $ld$ |
| $\eta_{re}$ | Repair Proteins | 6350 | aa/molecule of $re$ |
| $\eta_{in}$ | Internal Nitrogen | 14 | Da |
| $\eta_{ic}$ | Internal Carbon | 12 | Da |
| $\eta_{aa}$ | Amino Acid | 110 | Da |
| $\eta_{gu}$ | Glucose | 180 | Da |
| $\eta_{li}$ | Lipid Storage | 800 | Da |

| Space and density constraints |  |  |  |
| --- | --- | --- | --- |
| $s_{tr}$ | Transporter surface area | $1.26 \times 10^{-5}$ | $\mu m^2$ |
| $s_{lm}$ | Lipid surface area | $0.5 \times 10^{-6}$ | $\mu m^2$ |
| $M_{max}$ | Transporter to lipid ratio | Table S3 | unitless |
| $V_p$ | Photosystem volume | Table S3 | $\mu m^3$ |
| $V_{li}$ | Lipid volume | $1.5 \times 10^{-9}$ | $\mu m^3$ |
| $V_{nu}$ | Nucleus volume | Table S3 | $\mu m^3$ |
| $D_{max}$ | Maximum density | $1.8 \times 10^{10}$ | Da $\mu m^{-3}$ |
| $\alpha$ | Maximum antioxidant concentration | Table S3 | antioxidants $\mu m^{-3}$ |
| $\alpha_{min}$ | Minimum antioxidant concentration | 200 | antioxidants $\mu m^{-3}$ |
| Carbon and Nitrogen quotas |  |  |  |
| $\eta_{aac}$ | Amino Acids Carbon | 5 | C/aa |
| $\eta_{guc}$ | Glucose Carbon | 6 | C/molecule of *gu* |
| $\eta_{lic}$ | Lipids Carbon | 16 | C/molecule of *li* |
| $q_{pt}$ | Protein quota | 0.20 | N/C |
| $q_p$ | Photosystems quota | 0.10 | N/C |
| $q_{ri}$ | Ribosomes quota | 0.33 | N/C |
| Energy conversion factors |  |  |  |
| $e_p$ | Photosystems | 1.0 | ATP/photon |
| $e_{ru}$ | rubisco | 10 | ATP/C |
| $e_{tr}$ | Transporters | 1.0 | ATP/N |
| $e_{ri}$ | Protein synthesis | 48 | ATP/aa |
| $e_{gl}$ | Glycolysis | 5.0 | ATP/C |
| $e_{lb}$ | Lipid Synthesis | 3.9 | ATP/C |
| $e_{ld}$ | Lipid Degradation | 6.6 | ATP/C |
| $e_{re}$ | Repair Proteins | 4.5 | ATP/photosystem |
| $e_d$ | Heat damage | $2.0 \times 10^7/\alpha$ | ATP/photosystem |

**Table S3:** Parameter values for the different phytoplankton functional types. Parameter values used to parameterize the different phytoplankton functional types represented in our latitudinal simulations, i.e., *Prochlorococcus*, *Synechococcus*, warm-adapted diatom, and cold-adapted diatom. More details and sources can be found in the Supplementary Text.

| Parameter | Pro | Syn | W-Diatom | C-Diatom | Unit | Description |
| --- | --- | --- | --- | --- | --- | --- |
| $E_a$ | 0.85 | 1.2 | 1.2 | 0.65 | eV | Thermal dependency |
| $T_d$ | 35 | 35 | 35 | 25 | °C | Stress temperature |
| $K_{tr}$ | 0.1 | 2.0 | 60.0 | 60.0 | $\mu M$ | Half-saturation N-uptake |
| $V_p$ | $1.5e-5$ | $1.5e-5$ | $3e-5$ | $3e-5$ | $\mu m^3$ | Photosystem volume |
| $V_{nu}$ | 0.05 | 0.1 | 50 | 50 | $\mu m^3$ | Nucleus volume |
| $M_{max}$ | $1 \times 10^{-5}$ | $3 \times 10^{-5}$ | $4.5 \times 10^{-4}$ | $4.5 \times 10^{-4}$ | unitless | Transporter to lipid ratio |
| $\alpha$ | $8 \times 10^6$ | $1 \times 10^7$ | $2 \times 10^7$ | $2 \times 10^7$ | ant $\mu m^{-3}$ | Antioxidant concentration |

**Table S4:** Comparison between modeled and observed proteomic shifts in response to heat-stress. Thermal acclimation responses are shown where the arrows correspond to up-regulation (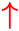) or down-regulation (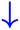) of pathways from populations growing at temperature optimum to populations subjected to heat-stress. Observations (proteomic data) for the enriched gene ontology (GO) terms correspond to the ancestral populations from an experimental evolution study with the marine diatom *Chaetoceros simplex* growing at 25 °C (optimum) versus the ancestral populations growing at 31 °C (heat-stress) or the warm-evolved populations growing at 34 °C (heat-stress)^52^.

| Model Variable | Model | GO terms | Ancestral | Evolved |
| --- | --- | --- | --- | --- |
| Transporters | ↓ | nitrate/nitrite transport | ↓ | ↓ |
|  |  | nitrate/nitrite transporter activity | ↓ | ↓ |
|  |  | nitrate reductase (NADPH) activity | not enriched | ↓ |
|  |  | nitrate assimilation | not enriched | ↓ |
| Glycolysis | (↑) | glycolytic process | (↑) | (↑) |
|  |  | fructose 1,6-bisphosphate metabolic process | (↑) | not enriched |
|  |  | phosphopyruvate hydratase activity | (↑) | not enriched |
|  |  | glucose 6-phosphate metabolic process | not enriched | (↑) |
| Heat-mitigation | (↑) | cell redox homeostasis | (↑) | (↑) |
|  |  | superoxide dismutase activity | (↑) | not enriched |
|  |  | removal of superoxide radicals | (↑) | not enriched |
|  |  | cellular response to oxygen compound | (↑) | not enriched |
| Repair proteins | (↑) | cellular response to heat | (↑) | (↑) |
|  |  | cellular response to unfolded protein | (↑) | (↑) |
|  |  | heat shock protein binding | (↑) | (↑) |
|  |  | protein folding chaperone | (↑) | (↑) |
|  |  | misfolded protein binding | (↑) | (↑) |
|  |  | positive regulation of DNA repair | (↑) | not enriched |
|  |  | unfolded protein binding | (↑) | (↑) |
|  |  | protein stabilization | not enriched | (↑) |

**Table S5:** Comparison between modeled (M) and observed (O) proteomic investment shifts in response to nutrient limitation. Acclimation responses are shown where the arrows correspond to the up-regulation (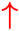) or down-regulation (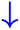) of pathways from nutrient-replete to nutrient-limiting or depleted conditions. Specific protein names used for the comparisons against each modeled pool are given in the Supplementary Methods.

| Functional Type | Variable | M | O | Reference |
| --- | --- | --- | --- | --- |
| Cyanobacteria | Transporters | (↑) | (↑) | 75 |
|  | Photosystems | ↓ | (↑) | 75 |
|  | Rubisco | ↓ | ↓ | 75 |
|  | Ribosomes | ↓ | ↓ | 75 |
|  | Glycolysis | ↓ | ↓ | 75 |
| Diatom | Transporters | (↑) | (↑) | 28,74,76 |
|  | Photosystems | ↓ | ↓ | 28,74,76 |
|  | Rubisco | ↓ | ↓ | 74,76 |
|  |  |  | (↑) | 28 |
|  | Ribosomes | ↓ | ↓ | 28,74,76 |
|  | Glycolysis | ↓ | ↓ | 28,76 |
|  |  |  | (↑) | 74 |
|  | Lipid synthesis | (↑) | (↑) | 28,76 |
|  |  |  | ↓ | 74 |

## References

1. IPCC (2023). Climate change 2023: Synthesis report. contribution of working groups i, ii and iii to the sixth assessment report of the intergovernmental panel on climate change. https://www.ipcc.ch/report/ar6/syr/.. Core Writing Team, H. Lee and J. Romero (eds.).

2. Kwiatkowski, L., Torres, O., Bopp, L., Aumont, O., Chamberlain, M., Christian, J.R., Dunne, J.P., Gehlen, M., Ilyina, T., John, J.G. et al. (2020). Twenty-first century ocean warming, acidification, deoxygenation, and upper-ocean nutrient and primary production decline from cmip6 model projections. Biogeosciences 17, 3439–3470.

3. Bopp, L., Monfray, P., Aumont, O., Dufresne, J.L., Le Treut, H., Madec, G., Terray, L., and Orr, J.C. (2001). Potential impact of climate change on marine export production. Global Biogeochemical Cycles 15, 81–99.

4. Boyce, D.G., Lewis, M.R., and Worm, B. (2010). Global phytoplankton decline over the past century. Nature 466, 591–596.

5. Thomas, M.K., Kremer, C.T., Klausmeier, C.A., and Litchman, E. (2012). A global pattern of thermal adaptation in marine phytoplankton. Science 338, 1085–1088.

6. Atkinson, A., Rossberg, A.G., Gaedke, U., Sprules, G., Heneghan, R.F., Batziakas, S., Grigoratou, M., Fileman, E., Schmidt, K., and Frangoulis, C. (2024). Steeper size spectra with decreasing phytoplankton biomass indicate strong trophic amplification and future fish declines. Nature Communications 15, 381.

7. Worden, A.Z., Follows, M.J., Giovannoni, S.J., Wilken, S., Zimmerman, A.E., and Keeling, P.J. (2015). Rethinking the marine carbon cycle: factoring in the multifarious lifestyles of microbes. Science 347, 1257594.

8. Raven, J.A., and Geider, R.J. (1988). Temperature and algal growth. New Phytol. 110, 441–461.

9. Boyd, P.W., Collins, S., Dupont, S., Fabricius, K., Gattuso, J.P., Havenhand, J., Hutchins, D.A., Riebesell, U., Rintoul, M.S., Vichi, M. et al. (2018). Experimental strategies to assess the biological ramifications of multiple drivers of global ocean change—a review. Global change biology 24, 2239–2261.

10. Van de Waal, D.B., and Litchman, E. (2020). Multiple global change stressor effects on phytoplankton nutrient acquisition in a future ocean. Philosophical Transactions of the Royal Society B 375, 20190706.

11. Litchman, E., Klausmeier, C., and Yoshiyama, K. (2009). Contrasting size evolution in marine and freshwater diatoms. Proceedings of the National Academy of Sciences 106, 2665–2670.

12. Finkel, Z.V., Beardall, J., Flynn, K.J., Quigg, A., Rees, T.A.V., and Raven, J.A. (2010). Phytoplankton in a changing world: cell size and elemental stoichiometry. Journal of plankton research 32, 119–137.

13. Marañón, E. (2015). Cell size as a key determinant of phytoplankton metabolism and community structure. Annual review of marine science 7, 241–264.

14. Sommer, U., Peter, K.H., Genitsaris, S., and Moustaka-Gouni, M. (2017). Do marine phytoplankton follow b ergmann’s rule sensu lato? Biological Reviews 92, 1011–1026.

15. Aksnes, D., and Egge, J. (1991). A theoretical model for nutrient uptake in phytoplankton. Marine ecology progress series. Oldendorf 70, 65–72.

16. Atkinson, D., Ciotti, B.J., and Montagnes, D.J. (2003). Protists decrease in size linearly with temperature: ca. 2.5% c-1. Proc. Royal Soc. B 270, 2605–2611.

17. Daufresne, M., Lengfellner, K., and Sommer, U. (2009). Global warming benefits the small in aquatic ecosystems. Proc. Natl. Acad. Sci. 106, 12788–12793.

18. Schaum, C.E., Buckling, A., Smirnoff, N., Studholme, D., and Yvon-Durocher, G. (2018). Environmental fluctuations accelerate molecular evolution of thermal tolerance in a marine diatom. Nature communications 9, 1719.

19. Liang, Y., Koester, J.A., Liefer, J.D., Irwin, A.J., and Finkel, Z.V. (2019). Molecular mechanisms of temperature acclimation and adaptation in marine diatoms. The ISME Journal 13, 2415–2425.

20. Barton, S., Padfield, D., Masterson, A., Buckling, A., Smirnoff, N., and Yvon-Durocher, G. (2023). Comparative experimental evolution reveals species-specific idiosyncrasies in marine phytoplankton adaptation to warming. Global Change Biology 29, 5261–5275.

21. Labban, A., Shibl, A.A., Calleja, M.L., Hong, P.Y., and Morán, X.A.G. (2023). Growth dynamics and transcriptional responses of a red sea prochlorococcus strain to varying temperatures. Environmental microbiology 25, 1007–1021.

22. Hattich, G., Jokinen, S., Sildever, S., Gareis, M., Heikkinen, J., Junghardt, N., Segovia, M., Machado, M., and Sjöqvist, C. (2024). Temperature optima of a natural diatom population increases as global warming proceeds. Nature Climate Change 14, 518–525.

23. Leles, S.G., and Levine, N.M. (2023). Mechanistic constraints on the trade-off between photosynthesis and respiration in response to warming. Science Advances 9, eadh8043.

24. Longhurst, A.R. (2010). Ecological geography of the sea. Elsevier.

25. Anderson, S.I., Fronda, C., Barton, A.D., Clayton, S., Rynearson, T.A., and Dutkiewicz, S. (2024). Phytoplankton thermal trait parameterization alters community structure and biogeochemical processes in a modeled ocean. Global Change Biology 30, e17093.

26. Dutkiewicz, S., Scott, J.R., and Follows, M. (2013). Winners and losers: Ecological and biogeochemical changes in a warming ocean. Global Biogeochemical Cycles 27, 463–477.

27. Flombaum, P., Gallegos, J.L., Gordillo, R.A., Rincón, J., Zabala, L.L., Jiao, N., Karl, D.M., Li, W.K., Lomas, M.W., Veneziano, D. et al. (2013). Present and future global distributions of the marine cyanobacteria prochlorococcus and synechococcus. Proceedings of the National Academy of Sciences 110, 9824–9829.

28. Yang, Z.K., Ma, Y.H., Zheng, J.W., Yang, W.D., Liu, J.S., and Li, H.Y. (2014). Proteomics to reveal metabolic network shifts towards lipid accumulation following nitrogen deprivation in the diatom phaeodactylum tricornutum. Journal of Applied Phycology 26, 73–82.

29. Baroni, É.G., Yap, K.Y., Webley, P.A., Scales, P.J., and Martin, G.J. (2019). The effect of nitrogen depletion on the cell size, shape, density and gravitational settling of nannochloropsis salina, chlorella sp.(marine) and haematococcus pluvialis. Algal Research 39, 101454.

30. Marañón, E., Fernández-González, C., and Tarran, G.A. (2024). Effect of temperature, nutrients and growth rate on picophytoplankton cell size across the atlantic ocean. Scientific Reports 14, 28034.

31. Miettinen, T.P., Gomez, A.L., Wu, Y., Wu, W., Usherwood, T.R., Hwang, Y., Roller, B.R., Polz, M.F., and Manalis, S.R. (2024). Cell size, density, and nutrient dependency of unicellular algal gravitational sinking velocities. Science Advances 10, eadn8356.

32. Ribalet, F., Dutkiewicz, S., Monier, E., and Armbrust, E.V. (2025). Future ocean warming may cause large reductions in prochlorococcus biomass and productivity. Nature microbiology pp. 1–12.

33. Moore, C., Mills, M., Arrigo, K., Berman-Frank, I., Bopp, L., Boyd, P., Galbraith, E., Geider, R., Guieu, C., Jaccard, S. et al. (2013). Processes and patterns of oceanic nutrient limitation. Nature geoscience 6, 701–710.

34. Kristiansen, S. (1983). The temperature optimum of the nitrate reductase assay for marine phytoplankton 1. Limnol. Oceanogr. 28, 776–780.

35. Berges, J.A., Varela, D.E., and Harrison, P.J. (2002). Effects of temperature on growth rate, cell composition and nitrogen metabolism in the marine diatom thalassiosira pseudonana (bacillariophyceae). Mar. Ecol. Prog. Ser. 225, 139–146.

36. Baker, K.G., Robinson, C.M., Radford, D.T., McInnes, A.S., Evenhuis, C., and Doblin, M.A. (2016). Thermal performance curves of functional traits aid understanding of thermally induced changes in diatom-mediated biogeochemical fluxes. Front. Mar. Sci. 3, 44.

37. Tomczyk, N., Rosemond, A., Kaz, A., and Benstead, J. (2022). Contrasting activation energies of respiration and nutrient uptake drive lower ecosystem-level uptake at higher temperatures. Biogeosci. pp. 1–21.

38. Hemme, D., Veyel, D., Mühlhaus, T., Sommer, F., Jüppner, J., Unger, A.K., Sandmann, M., Fehrle, I., Schönfelder, S., Steup, M., et al. (2014). Systems-wide analysis of acclimation responses to long-term heat stress and recovery in the photosynthetic model organism chlamydomonas reinhardtii. Plant Cell 26, 4270–4297.

39. Jarc, E., and Petan, T. (2019). Focus: Organelles: Lipid droplets and the management of cellular stress. Yale J. Biol. Med. 92, 435.

40. McCain, J.S.P., and Bertrand, E.M. (2022). Phytoplankton antioxidant systems and their contributions to cellular elemental stoichiometry. Limnology and Oceanography Letters 7, 96–111.

41. Thomas, M.K., Aranguren-Gassis, M., Kremer, C.T., Gould, M.R., Anderson, K., Klausmeier, C.A., and Litchman, E. (2017). Temperature–nutrient interactions exacerbate sensitivity to warming in phytoplankton. Global change biology 23, 3269–3280.

42. Bestion, E., Schaum, C.E., and Yvon-Durocher, G. (2018). Nutrient limitation constrains thermal tolerance in freshwater phytoplankton. Limnology and Oceanography Letters 3, 436–443.

43. Aranguren-Gassis, M., and Litchman, E. (2020). Thermal performance of marine diatoms under contrasting nitrate availability. Journal of Plankton Research 42, 680–688.

44. Huey, R.B., and Kingsolver, J.G. (2019). Climate warming, resource availability, and the metabolic meltdown of ectotherms. The American Naturalist 194, E140–E150.

45. Edwards, K.F., Thomas, M.K., Klausmeier, C.A., and Litchman, E. (2012). Allometric scaling and taxonomic variation in nutrient utilization traits and maximum growth rate of phytoplankton. Limnology and Oceanography 57, 554–566.

46. Drábková, M., Admiraal, W., and Maršálek, B. (2007). Combined exposure to hydrogen peroxide and light selective effects on cyanobacteria, green algae, and diatoms. Environmental science & technology 41, 309–314.

47. Bernroitner, M., Zamocky, M., Furtmüller, P.G., Peschek, G.A., and Obinger, C. (2009). Occurrence, phylogeny, structure, and function of catalases and peroxidases in cyanobacteria. Journal of experimental botany 60, 423–440.

48. Rosenwasser, S., Graff van Creveld, S., Schatz, D., Malitsky, S., Tzfadia, O., Aharoni, A., Levin, Y., Gabashvili, A., Feldmesser, E., and Vardi, A. (2014). Mapping the diatom redox-sensitive proteome provides insight into response to nitrogen stress in the marine environment. Proceedings of the National Academy of Sciences 111, 2740–2745.

49. Mella-Flores, D., Six, C., Ratin, M., Partensky, F., Boutte, C., Le Corguillé, G., Marie, D., Blot, N., Gourvil, P., Kolowrat, C., et al. (2012). Prochlorococcus and synechococcus have evolved different adaptive mechanisms to cope with light and uv stress. Frontiers in microbiology 3, 285.

50. Anderson, S., Barton, A., Clayton, S., Dutkiewicz, S., and Rynearson, T. (2021). Marine phytoplankton functional types exhibit diverse responses to thermal change. Nature communications 12, 6413.

51. Danabasoglu, G., Lamarque, J.F., Bacmeister, J., Bailey, D., DuVivier, A., Edwards, J., Emmons, L., Fasullo, J., Garcia, R., Gettelman, A. et al. (2020). The community earth system model version 2 (cesm2). Journal of Advances in Modeling Earth Systems 12, e2019MS001916.

52. Aranguren-Gassis, M., Diz, A.P., Huete-Ortega, M., Allen, A., and Litchman, E. (2024). Evolution of thermal tolerance in marine diatoms: Metabolic strategies under heat stress. bioRxiv pp. 2024–11.

53. Duerschlag, J., Mohr, W., Ferdelman, T.G., LaRoche, J., Desai, D., Croot, P.L., Voß, D., Zielinski, O., Lavik, G., Littmann, S. et al. (2022). Niche partitioning by photosynthetic plankton as a driver of co2-fixation across the oligotrophic south pacific subtropical ocean. The ISME Journal 16, 465–476.

54. Chaidez, V., Dreano, D., Agusti, S., Duarte, C., and Hoteit, I. (2017). Decadal trends in red sea maximum surface temperature, sci. rep., 7, 8144..

55. Al-Otaibi, N., Huete-Stauffer, T.M., Calleja, M.L., Irigoien, X., and Morán, X.A.G. (2020). Seasonal variability and vertical distribution of autotrophic and heterotrophic picoplankton in the central red sea. PeerJ 8, e8612.

56. Coello-Camba, A., and Agustí, S. (2021). Picophytoplankton niche partitioning in the warmest oligotrophic sea. Frontiers in Marine Science 8, 651877.

57. Kheireddine, M., Ouhssain, M., Claustre, H., Uitz, J., Gentili, B., and Jones, B.H. (2017). Assessing pigment-based phytoplankton community distributions in the red sea. Frontiers in Marine Science 4, 132.

58. Morris, J.J., Lenski, R.E., and Zinser, E.R. (2012). The black queen hypothesis: evolution of dependencies through adaptive gene loss. MBio 3, 10–1128.

59. Weenink, E.F., Matthijs, H.C., Schuurmans, J.M., Piel, T., van Herk, M.J., Sigon, C.A., Visser, P.M., and Huisman, J. (2021). Interspecific protection against oxidative stress: green algae protect harmful cyanobacteria against hydrogen peroxide. Environmental Microbiology 23, 2404–2419.

60. Hennon, G.M., Morris, J.J., Haley, S.T., Zinser, E.R., Durrant, A.R., Entwistle, E., Dokland, T., and Dyhrman, S.T. (2018). The impact of elevated co2 on prochlorococcus and microbial interactions with ‘helper’bacterium alteromonas. The ISME journal 12, 520–531.

61. Lu, Z., Entwistle, E., Kuhl, M.D., Durrant, A.R., Filho, M.M.B., Goswami, A., and Morris, J. (2024). Marine phytoplankton and heterotrophic bacteria rapidly adapt to future pco2 conditions in experimental co-cultures. bioRxiv pp. 2024–02.

62. Schvarcz, C.R., Wilson, S.T., Caffin, M., Stancheva, R., Li, Q., Turk-Kubo, K.A., White, A.E., Karl, D.M., Zehr, J.P., and Steward, G.F. (2022). Overlooked and widespread pennate diatom-diazotroph symbioses in the sea. Nature communications 13, 799.

63. Coale, T.H., Loconte, V., Turk-Kubo, K.A., Vanslembrouck, B., Mak, W.K.E., Cheung, S., Ekman, A., Chen, J.H., Hagino, K., Takano, Y. et al. (2024). Nitrogen-fixing organelle in a marine alga. Science 384, 217–222.

64. Amin, S.A., Parker, M.S., and Armbrust, E.V. (2012). Interactions between diatoms and bacteria. Microbiology and Molecular Biology Reviews 76, 667–684.

65. Ward, B.A., and Follows, M.J. (2016). Marine mixotrophy increases trophic transfer efficiency, mean organism size, and vertical carbon flux. Proceedings of the National Academy of Sciences 113, 2958–2963.

66. Bezanson, J., Edelman, A., Karpinski, S., and Shah, V.B. (2017). Julia: A fresh approach to numerical computing. SIAM review 59, 65–98. URL: 10.1137/141000671.

67. Mizrachi, A., Graff van Creveld, S., Shapiro, O.H., Rosenwasser, S., and Vardi, A. (2019). Light-dependent single-cell heterogeneity in the chloroplast redox state regulates cell fate in a marine diatom. Elife 8, e47732.

68. Alonso-Sáez, L., Palacio, A.S., Cabello, A.M., Robaina-Estévez, S., González, J.M., Gar-czarek, L., and López-Urrutia, Á. (2023). Transcriptional mechanisms of thermal acclimation in prochlorococcus. MBio 14, e03425–22.

69. Dutkiewicz, S., Boyd, P.W., and Riebesell, U. (2021). Exploring biogeochemical and ecological redundancy in phytoplankton communities in the global ocean. Global change biology 27, 1196–1213.

70. Arshad, R., Calvaruso, C., Boekema, E.J., Büchel, C., and Kouřil, R. (2021). Revealing the architecture of the photosynthetic apparatus in the diatom thalassiosira pseudonana. Plant Physiol. 186, 2124–2136.

71. Malerba, M.E., and Marshall, D.J. (2021). Larger cells have relatively smaller nuclei across the tree of life. Evolution Letters 5, 306–314.

72. Finkel, Z.V., Follows, M.J., Liefer, J.D., Brown, C.M., Benner, I., and Irwin, A.J. (2016). Phylogenetic diversity in the macromolecular composition of microalgae. PLoS One 11, e0155977.

73. Aranguren-Gassis, M., Kremer, C.T., Klausmeier, C.A., and Litchman, E. (2019). Nitrogen limitation inhibits marine diatom adaptation to high temperatures. Ecology letters 22, 1860–1869.

74. Remmers, I.M., D’Adamo, S., Martens, D.E., de Vos, R.C., Mumm, R., America, A.H., Cordewener, J.H., Bakker, L.V., Peters, S.A., Wijffels, R.H., et al. (2018). Orchestration of transcriptome, proteome and metabolome in the diatom phaeodactylum tricornutum during nitrogen limitation. Algal research 35, 33–49.

75. Domínguez-Martín, M.A., Gómez-Baena, G., Díez, J., López-Grueso, M.J., Beynon, R.J., and García-Fernández, J.M. (2017). Quantitative proteomics shows extensive remodeling induced by nitrogen limitation in prochlorococcus marinus ss120. MSystems 2, 10–1128.

76. Chen, X.H., Li, Y.Y., Zhang, H., Liu, J.L., Xie, Z.X., Lin, L., and Wang, D.Z. (2018). Quantitative proteomics reveals common and specific responses of a marine diatom thalassiosira pseudonana to different macronutrient deficiencies. Frontiers in Microbiology 9, 2761.

